# SMB controls decompartmentalization in Arabidopsis root cap cells to execute programmed cell death

**DOI:** 10.1101/2023.08.10.552584

**Authors:** Jie Wang, Norbert Bollier, Rafael Andrade Buono, Hannah Vahldick, Zongcheng Lin, Qiangnan Feng, Roman Hudecek, Qihang Jiang, Evelien Mylle, Daniel Van Damme, Moritz K. Nowack

## Abstract

Programmed cell death (PCD) is a fundamental cellular process crucial to development, homeostasis, and immunity in multicellular eukaryotes. In contrast to our knowledge on the regulation of diverse animal cell death subroutines, information on execution of PCD in plants remains fragmentary. Here we make use of the accessibility of the *Arabidopsis thaliana* root cap to visualize the execution process of developmentally controlled PCD. We identify a succession of selective decompartmentalization events and ion fluxes that are controlled by a gene regulatory network downstream of the NAC transcription factor SOMBRERO (SMB). Surprisingly, breakdown of the large central vacuole is a relatively late and variable event, preceded by an increase of intracellular calcium levels and acidification, release of mitochondrial matrix proteins, leakage of nuclear and endoplasmic reticulum lumina, and release of fluorescent membrane reporters into the cytosol. Elevated intracellular calcium levels and acidification are sufficient to trigger cell death execution specifically in cells that are rendered competent to undergo PCD by SMB activity, suggesting that these ion fluxes act as PCD-triggering signals.

## Introduction

Programmed cell death (PCD) is a fundamental cellular process that fulfills a plethora of vital functions in multicellular organisms. In plants, PCD is both part of the response to biotic and abiotic environmental stresses, as well as an indispensable part of regular development (Huysmans et al., 2017; Kabbage et al., 2017). Developmentally controlled PCD (dPCD) occurs during vegetative and reproductive development, and is crucial for varied processes such as pollen formation, seed development, formation of the water-conducting xylem, or root cap turnover (Daneva et al., 2016). In many cases dPCD functions as the ultimate differentiation step of specific cell types, ending the vital functions and removing unwanted or no longer needed cells, or creating functional tissues composed of modified cell corpses (Daneva et al., 2016; Huysmans et al., 2017). Despite the importance of cell death processes during plant development, there is only little mechanistic knowledge on the regulation and execution of dPCD in plants.

As an integral part of cellular differentiation, dPCD has been divided into different phases, including preparation, triggering, execution, and post-mortem cell clearance (Van Durme and Nowack, 2016; Huysmans et al., 2017). During dPCD preparation, the transcriptional activation of dPCD-associated genes by transcription factors has been linked to the acquisition of dPCD competence (Olvera-Carrillo et al., 2015; Jiang et al., 2021). While mutation or misexpression of key transcription factors has demonstrated that transcriptional preparation of cell death is both necessary and sufficient for dPCD execution (Fendrych et al., 2014; Huysmans et al., 2018; Cubría-Radío and Nowack, 2019), the function of all but a few downstream target genes, and therefore the actual dPCD execution mechanism, remains unclear.

Over the last years, the Arabidopsis root cap has emerged as a powerful model system to study dPCD in its native developmental context *in planta*. The root cap is located at the tip of the growing plant root, ensheathing the root apical meristem. As the other cells in the root meristem, the root cap is generated by specific stem cells proximal to the quiescent center that iteratively produce new layers of columella and lateral root cap (LRC) cells. While the columella differentiates into gravity-sensing cells at the very root tip, LRC cells divide and flank the meristem up to the start of the elongation zone (Kumpf and Nowack, 2015). In this zone, epidermis cells start elongating while the overlying LRC cells undergo dPCD followed by a complete corpse clearance on the root surface that rapidly removes any trace of dead LRC cells (Fendrych et al., 2014).

The root cap-specific NAC (No apical meristem, Arabidopsis thaliana activating factor, Cup-shaped cotyledon) transcription factor SOMBRERO (ANAC033/SMB) controls key steps of root cap development, including the exit from root cap stem cell fate (Willemsen et al., 2008), root cap maturation (Bennett et al., 2010), and finally dPCD execution and post-mortem corpse clearance in concert with ANAC087 and ANAC046 (Fendrych et al., 2014; Huysmans et al., 2018). In *smb-3* mutants, cell death is delayed and occurs in the root elongation zone, canonical dPCD-associated genes are not expressed, and postmortem corpse clearance does not occur (Fendrych et al., 2014; Olvera-Carrillo et al., 2015; Huysmans et al., 2018). Interestingly, LRC cell death in the *smb-3* mutant is suppressed upon pharmacological reduction of root elongation, suggesting that the delayed and aberrant cell death *smb-3* result is a non-controlled result of physical stresses (Fendrych et al., 2014).

How dPCD execution in plants is accomplished on a cellular and mechanistic level remains little understood. Many dPCD processes have been characterized as forms of “vacuolar PCD”, in which a gradual volume increase of the central vacuole precedes the rupture of the vacuolar membrane (tonoplast). Mitochondria and other organelles, including the plasma membrane, remain intact until tonoplast rupture (van Doorn et al., 2011). Vacuolar collapse is thought to release a cocktail of vacuolar proteins including hydrolytic enzymes to accomplish both cell death execution and partial or complete post-mortem corpse clearance (Hara-Nishimura and Hatsugai, 2011). However, there exists little information on the exact sequence of subcellular events of dPCD execution.

The situation of the Arabidopsis root cap on the root periphery makes it a convenient dPCD model system to analyze the cellular processes occurring during dPCD execution in a high temporal resolution in its endogenous developmental context. Previously, we had described plasma membrane (PM) permeation for propidium iodide (PI) and vacuolar collapse to occur after a pronounced intracellular acidification (Fendrych, 2014). Here we present a systematic multifactor time-course analysis of the cellular decompartmentalization processes that occur during dPCD execution. By establishing nuclear envelope breakdown as a reference time point, we reveal that disintegration of mitochondria and the endoplasmic reticulum (ER) occur early, followed only later by vacuolar collapse and PM permeation for PI. Simultaneously with NE breakdown, the cytosolic leaflet of the PM sheds membrane-bound reporter proteins, suggesting a profound modification of the membrane proteome. Surprisingly, we found that PM permeation is only partial; the PM remains impermeable for reporter proteins during post-mortem corpse clearance. Preceding these cellular changes, we detected intracellular acidification coupled with a transient increase of intracellular calcium levels. These events and their order of appearance are highly aberrant in *smb-3* mutants, demonstrating that they are part of a genetically controlled dPCD execution program orchestrated downstream of SMB. Accordingly, pharmacologically lowering intracellular pH and increasing calcium levels is sufficient to trigger dPCD execution in dPCD-competent root cap cells in the wild type, but not in the *smb-3* mutant.

## Results

### dPCD execution follows a sequence of successive membrane permeation events

The distalmost LRC cells continuously undergo dPCD (Figure 1A) providing an ideal model to investigate the cellular characteristics of dPCD execution. During dPCD execution, vacuolar collapse and loss of plasma membrane (PM) integrity indicated by PI entry have been described. However, both events occur in a widely variable time frame after intracellular acidification, which has been observed in several plant dPCD contexts (Young et al., 2010; Fendrych et al., 2014; Wilkins et al., 2015).

**Figure 1:**
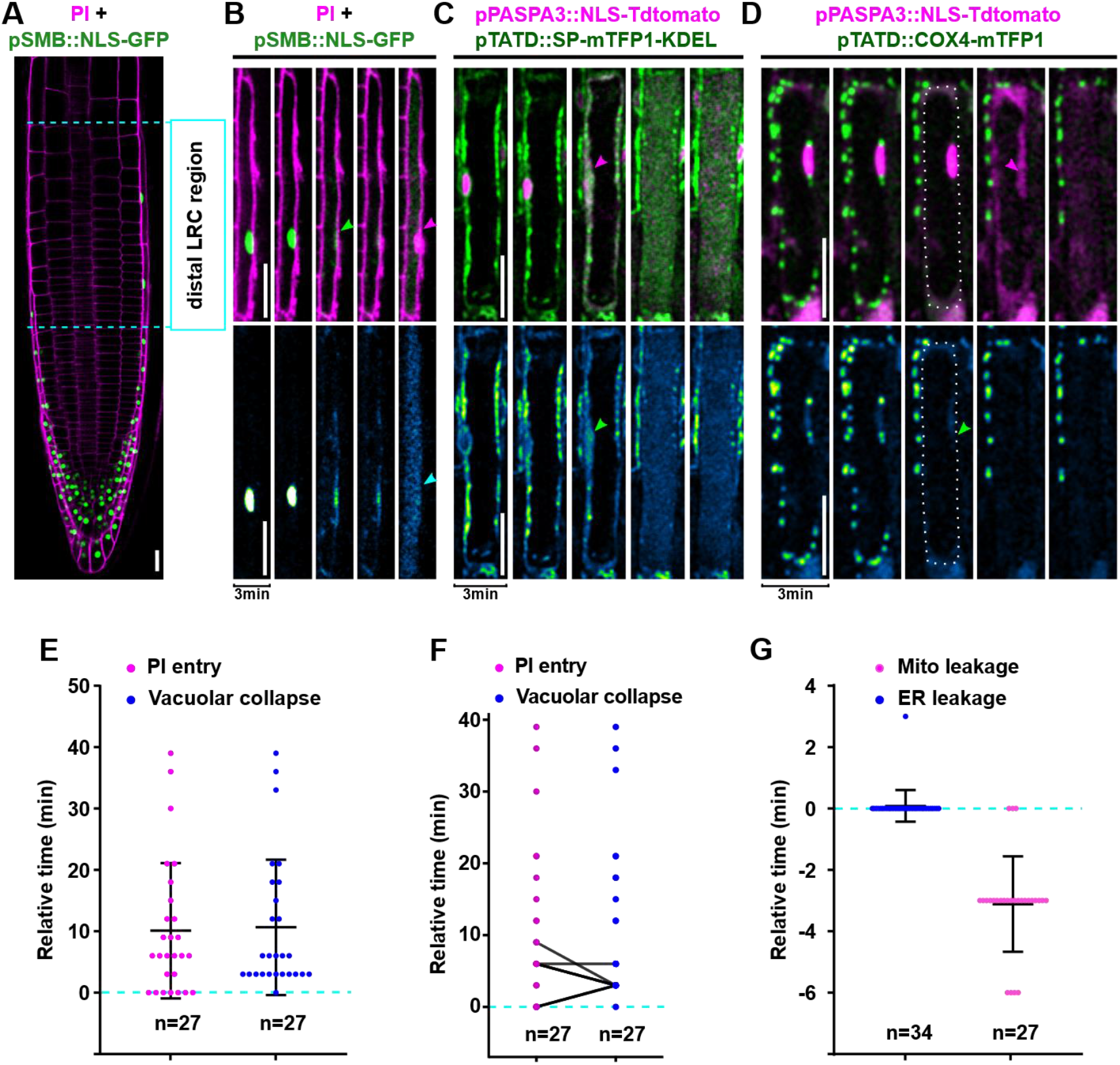
PCD execution occurs in a sequence of successive cellular decompartmentalization events. **(A)** Confocal image of a root tip from a 4-day-old seedling expressing nuclear reporter *pSMB::NLS-GFP* stained with PI. The distal LRC region in which cells prepare for and execute PCD is highlighted by dashed lines. **(B-D)** Confocal time-lapse series of dying LRC cells. Upper panels show PI or NLS-TdTOMATO in magenta and GFP or mTFP1 in green, the lower panels show the GFP or mTFP1 channel separately. **(B)** A cell expressing *pSMB::NLS-GFP* stained with PI. Nuclear envelope (NE) breakdown (green arrow) is followed by PI entry (magenta arrow) and vacuolar collapse (cyan arrow). **(C)** A cell expressing *pPASPA3::NLS-TdTOMATO* and *pTATD::SP-mTFP1-KDEL*. Magenta arrow marks NE breakdown, green arrow indicates ER leakage into the cytoplasm. **(D)** A cell expressing *pPASPA3::NLS-TdTOMATO* and *pTATD::COX4-mTFP1*. The dashed line marks the contour of the dying LRC cell, the green arrow indicates mitochondria leakage followed by NE breakdown (magenta arrow). **(E-G)** Quantification of decompartmentalization timing in relation to NE breakdown (dashed line); indicated is the mean ± standard deviation, n indicates the number of dying cells analyzed in at least 5 different roots each. **(E)** Timing of PI entry and vacuolar collapse, **(F)** correlation of PI entry and vacuolar collapse in individual cells connected by lines, **(G)** timing of ER and mitochondria leakage. Scale bars are 20 μm.

To establish an easy-to-detect reference time point of dPCD execution, we chose to visualize the death-associated disruption of the nucleocytoplasmic barrier (Domínguez and Cejudo, 2012) as a release of nuclear-localized fluorescent proteins (NLS-FPs) into the cytosol. Live-cell time-lapse imaging of NLS-GFP reporter lines in 3-minute intervals revealed an abrupt decrease of nuclear fluorescent signal caused by the diffusion of NLS-GFP into the cytosol (Figure 1B, Movie1). Release of NLS-FPs into the cytosol generates a binary readout that we utilized as a reference time point to calibrate other cellular processes occurring during dPCD execution.

NE permeation occurs prior to vacuolar collapse, which is visualized by a uniform distribution of fluorescent protein throughout the cell volume (Figure 1B-C). Combined with PI staining, we revealed that both vacuolar collapse and PM permeabilization for PI occur on average around 10 minutes after NE permeation, though both events happen remarkably variably and as late as 40 minutes after NE breakdown (Figure 1 B, E). Interestingly, PI entry and vacuolar collapse are temporally correlated with each other in most cell death events (Figure 1F).

Next, we followed the release of ER lumen proteins into the cytosol and nucleoplasm using an ER-localized SP-mTFP1-KDEL reporter in combination with a nuclear-localized NLS-tdTOMATO. This pH-stable, dual color reporter combination revealed that ER leakage occurred simultaneously with NE leakage, as indicated by a mixing of mTFP1 and tdTOMATO signals (Figure 1C, G, Movie2). This finding suggests a tightly correlated permeation of these closely associated membrane systems.

Finally, we visualized mitochondrial integrity by imaging a mitochondrial matrix marker COX4-mTFP1 (Nelson et al., 2007) combined with the NLS-tdTOMATO nuclear reporter. We observed numerous small foci in the cytoplasm, revealing the abundance of small, discrete mitochondria typical for plant cells (Logan, 2006). The foci abruptly disappeared during dPCD execution, suggesting a release of mitochondrial matrix proteins into the cytosol (Figure 1D, Movie3). Interestingly, mitochondrial disintegration occurred on average 3 minutes before NE and ER leakage (Figure 1G).

Our findings indicate that cellular decompartmentalization during dPCD in root cap cells is a highly ordered decompartmentalization process that starts with the mitochondria, continues with NE and ER leakage, while tonoplast rupture and PM permeabilization for PI variably occur at later time points.

### dPCD execution modifies the PM, but leaves it impermeable for proteins

The PM constitutes a selective barrier between the cell and its environment that is crucial for the vital functions of a cell. Loss of PM integrity as visualized by entry of non-membrane permeable dyes has been used to detect cell death in animals and plants (Truernit and Haseloff, 2008; Rieger et al., 2010). To investigate PM properties during dPCD execution in the root cap, we imaged LRC cells expressing NLS-GFP under the root cap-specific *pSMB* promoter stained with a short pulse of the styryl dye FM4-64. Our data reveals that the PM remains present after NE leakage and vacuolar collapse (Figure 2A). Next, we used a pH-stable PM marker, CPK17-mTFP1, to visualize the PM during dPCD execution. Surprisingly, the mTFP1 signal dissociated from the PM during dPCD execution and became first soluble in the cytosol, then upon vacuolar collapse uniformly filled the entire cell while the PM remained present as indicated by FM4-64 staining (Figure 2B and Figure S1B). Imaging a co-expressed nuclear and membrane reporter revealed that shedding of the PM reporter occurred simultaneously with NE leakage (n= 26 cells, Figure 2C and Movie4). Both mTFP1 and tdTOMATO signals were detected inside the cell corpse, but not in the apoplast, even after the PM became permeable for PI (Figure 2B-C, Figure S1B-C). Moreover, we observed the formation of membrane-bound extracellular vesicles (EVs) when imaging pPASPA3::ToIM, a double-reporter combining cytosolic GFP and vacuolar localized RFP (Fendrych et al., 2014). Time-lapse imaging revealed the formation of EVs filled with fluorescent proteins remained intact for several hours after cell death execution (Figure 2 E and Movie5). Possibly, these EVs are blebbing out of cell corpses in the root elongation zone where LRC cell corpses are mechanically stretched. Regardless of their actual mode of formation, these EVs confirm that fluorescent proteins do not dissipate out of membrane-bound cell corpses, suggesting that PM-derived membranes remain impermeable for proteins of similar sizes during post-mortem corpse clearance.

**Figure 2:**
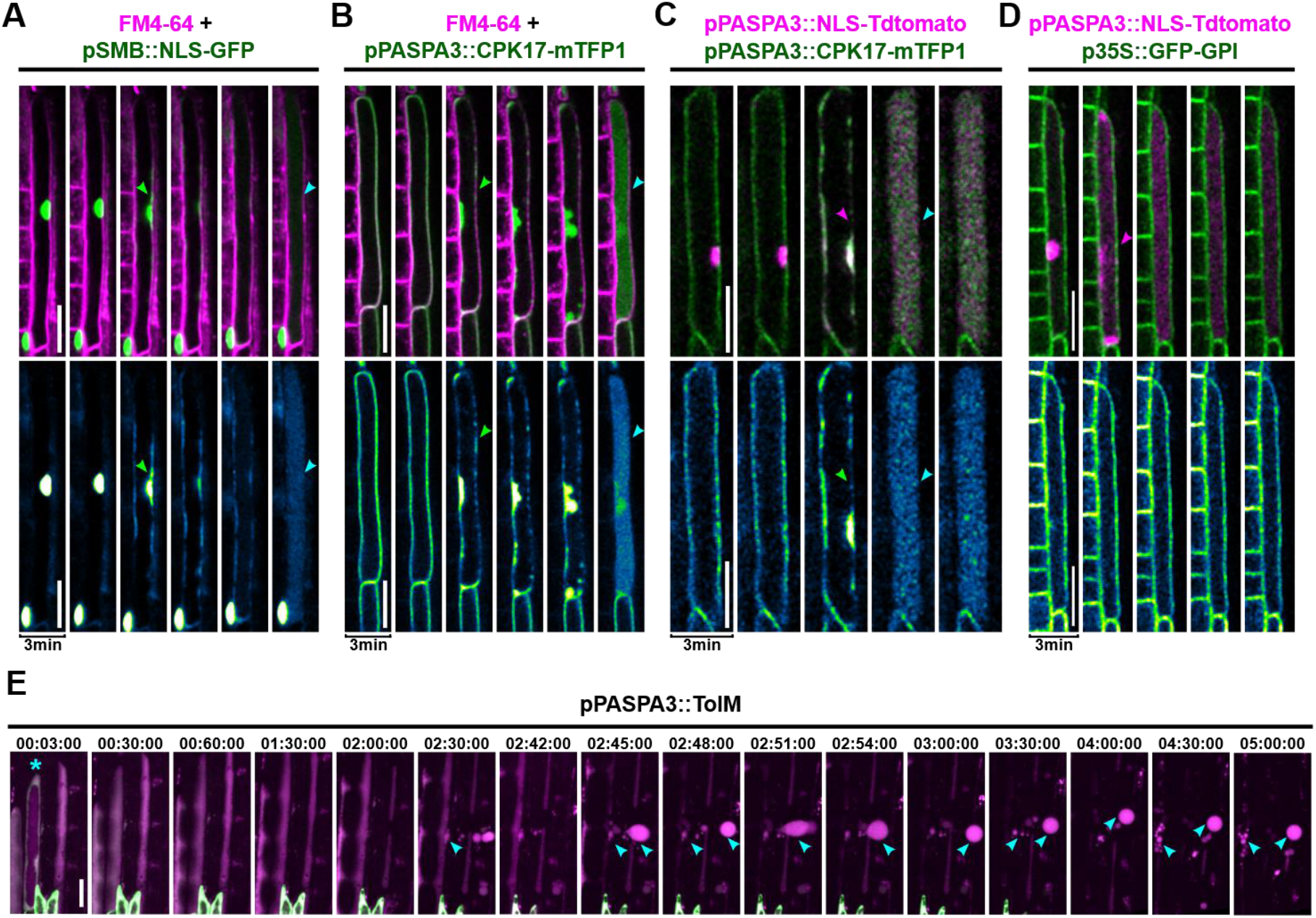
The plasma membrane remains impermeable for fluorescent proteins during and after PCD execution. **(A-D)** Confocal time-lapse series of dying LRC cells. Upper panels show FM4-64 or NLS-TdTOMATO in magenta and GFP or mTFP1 in green, the lower panels show the GFP or mTFP1 channel separately. **(A)** The PM stained by FM4-64 remains intact (cyan arrow) after NE breakdown (green arrow). **(B)** The PM reporter *pPASPA::CPK17-mTFP1* is shed off the PM (green arrow) while the PM remains intact after vacuolar collapse (cyan arrow). **(C)** Shedding of CPK17-mTFP1 from the PM and NE breakdown occur simultaneously (magenta arrow) prior to vacuolar collapse (cyan arrow), n=26 dying cells from 11 individual roots. **(D)** An apoplastic *p35S::GFP-GPI* PM marker remains on the PM after NE breakdown and vacuolar collapse (magenta arrow), n=8 roots were imaged. **(E)** Confocal time-lapse series of dying LRC cells expressing *pPASPA3::ToIM*. The cyan asterisk marks a cell prior to vacuolar collapse, cyan arrows indicate the generation of “extracellular vesicles” after cell death execution. Scale bars are 20 μm.

Nevertheless, the PM is subjected to modifications during dPCD execution, as indicated by its permeabilization for PI and the solubilization of CPK17-mTFP1. We confirmed solubilization of membrane-localized fluorescent proteins by imaging a line expressing a PM marker construct, pUBQ10::3xmCHERRY-SYP122 (Figure S1A, D). In addition to CPK17 and SYP122, which are predicted to contain one transmembrane domain, SCAMP5-GFP, a PM-localized reporter with 4 predicted transmembrane domains (Yperman et al., 2021), and the tonoplast reporter VAMP711-YFP (Feng et al., 2017) showed the same solubilization pattern during dPCD execution (Figure S1E-F). A shared commonality between these membrane markers is that the fluorescent moiety is predicted to be localized on the cytoplasmic side of the membrane. To investigate whether solubilization also affects fluorescent moieties localized on the apoplastic side of the PM, we investigated a GPI-anchored GFP marker (Martiniere et al., 2012). Intriguingly, time-lapse imaging revealed that GFP-GPI remains attached to the PM throughout dPCD execution (Figure 2D and Figure S1G). Imaging a dual marker line expressing 3xmCHERRY-SYP122 and GFP-GPI confirmed that the solubilization exclusively affects the fluorescent moieties on the cytoplasmic side of the PM (Figure S1H).

In sum, these data suggest that the PM undergoes specific modifications that are characterized by the solubilization of cytoplasmic membrane domains and by a partial loss of membrane impermeability for small charged molecules. However, while the PM does become permeable for PI, it does not disintegrate and remains impermeable for proteins the size of fluorescent reporters during post-mortem corpse clearance.

### *smb-3* undergoes disordered and delayed cell death execution

To evaluate which of the observed membrane modification processes are genetically controlled by the key root cap dPCD regulator SMB, we analyzed cell death processes in the *smb-3* mutant (Willemsen et al., 2008). Analysis of an *smb-3* line carrying *pSMB::NLS-YFP* (Kamiya et al., 2016) revealed a gradual fading of nuclear YFP fluorescence in dying cells (Figure 3A and D). This observation suggests that either acidification or NE permeation, or both, occurred in a more gradual fashion than during wild-type dPCD (Figure 3A, Movie6). Staining with PI showed a variable but slightly earlier PM permeabilization for PI in relation to NE leakage (Figure 3A). Also, the timing of vacuolar collapse was divergent in *smb-3*, occurring earlier than in the wild type (Figure 3B and D).

**Figure 3:**
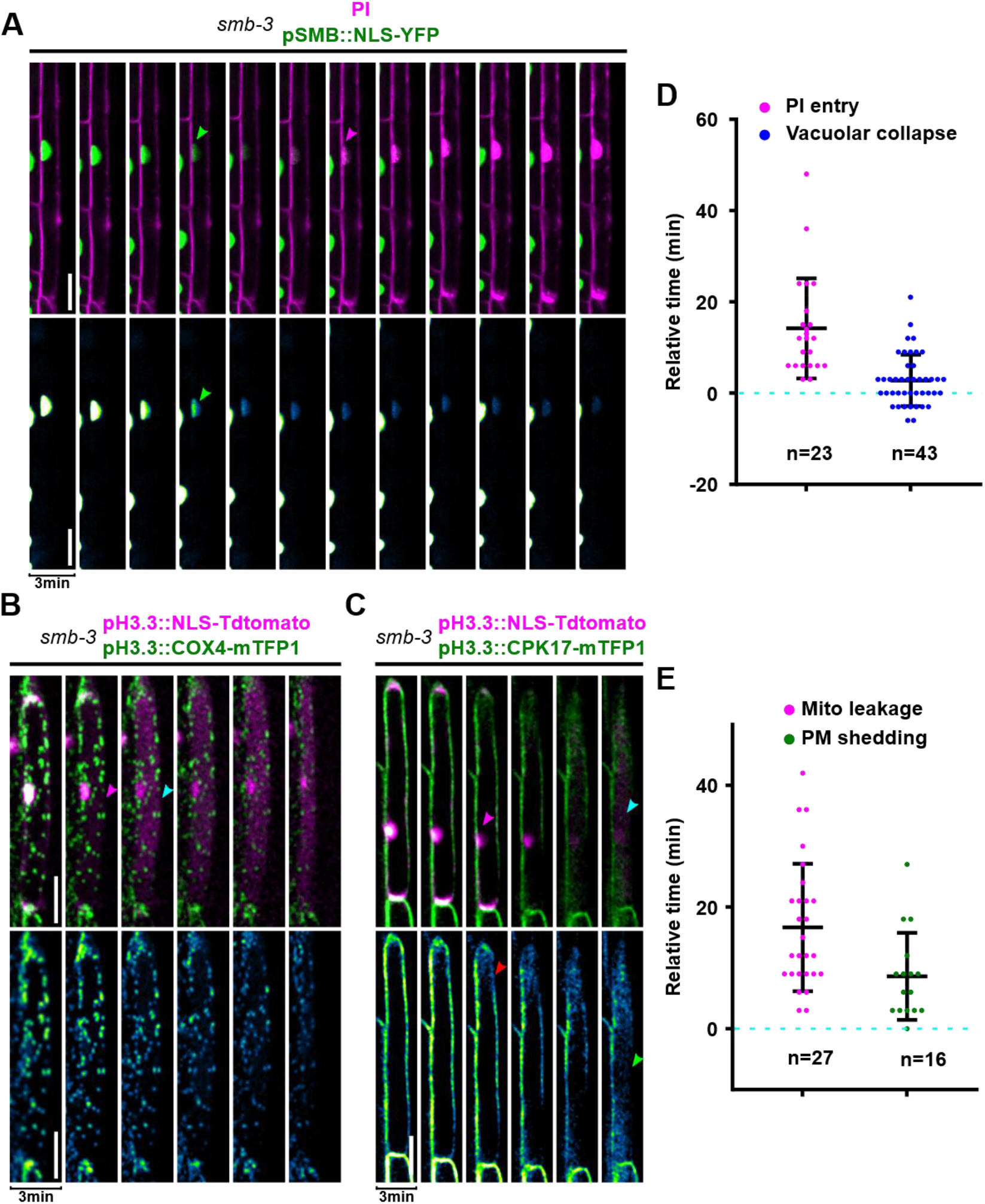
Aberrant decompartmentalization during cell death in the *smb-3* mutant. **(A-C)** Confocal time-lapse series of dying *smb-3* mutant LRC cells. Upper panels show PI or NLS-TdTOMATO in magenta and GFP or mTFP1 in green, the lower panels show the GFP or mTFP1 channel separately. **(A)** NE breakdown (green arrow) occurs more gradually than in the wild type, though still before PI entry (magenta arrow). **(B)** Mitochondria remain intact after NE breakdown (magenta arrow) and vacuolar collapse (cyan arrow), and mitochondrial signals only gradually fade away. **(C)** Shedding of a CPK17-mTFP1 PM reporter occurs partially before (red arrow) and is completed (green arrow) after NE breakdown occurs (magenta arrow) in *smb-3* mutants. **(D-E)** Quantification of decompartmentalization timing in relation to NE breakdown (dashed line) in dying *smb-3* mutant LRC cells; indicated is the mean ± standard deviation, n indicates the number of dying cells analyzed in at least 6 different roots each. **(D)** Timing of PI entry and vacuolar collapse, **(E)** timing of fading of mitochondrial signals and completion of mTFP1 shedding from the PM. Scale bars are 20 μm.

Analysis of a dual nuclear and ER-lumen reporter in the *smb-3* mutant background confirmed that NE breakdown occurred more gradually than in the wild type (Figure S2). Furthermore, no clear release of ER-resident fluorescent proteins could be observed, rather a gradual fading of the fluorescent signal that still localized to concrete foci (Figure S2). While this behavior renders the quantification of ER leakage timing impracticable, it suggests that ER leakage in *smb-3* does not occur in the same ordered fashion as in the wild type.

Analysis of mitochondrial integrity in the *smb-3* mutant revealed that the leakage of mitochondria was strongly delayed or even entirely absent, as COX4-mTFP1 foci remained visible after NE leakage and only gradually faded away (Figure 3B and E, Movie7). Finally, CPK17-mTFP1 time-lapse imaging revealed that PM-reporter solubilization did not occur simultaneously with NE leakage as in the wild type, but rather in a delayed and more gradual fashion (Figure 3C and E, Movie8). In contrast to the wild type, we could not observe a clear cytosolic mTFP1 signal during cell death in *smb-3* (Figure 3C).

Taken together these observations suggest that the order and the accuracy of cellular decompartmentalization is actively controlled by a genetic pathway downstream of the key regulator SMB. This implies that still unknown direct or indirect targets of SMB are responsible for the coordination and efficient execution of NE and ER leakage, mitochondrial disintegration, cytosolic membrane-reporter solubilization, and vacuolar collapse.

### Intracellular calcium elevation and acidification are aberrant in *smb-3* mutants

Plant cell death has been correlated with intracellular acidification and increased cytosolic calcium ion levels (Fendrych et al., 2014; Lin et al., 2020). We monitored cellular Ca^2+^ signatures by means of the genetically encoded biosensor Yellow Chameleon (YC3.6) (Nagai et al., 2004; Krebs et al., 2012; Lin et al., 2020). Targeting YC3.6 to the nucleus enabled us to directly correlate Ca^2+^ dynamics with NE leakage. Analysis of the dynamic relative [Ca^2+^]_nuc_ levels during dPCD execution revealed a sharp [Ca^2+^]_nuc_ transient during dPCD execution, which most often was recorded 3 minutes before NE breakdown (19/26 cells, Figure 4A and C-E, Figure S3A, Movie9). Next, we monitored relative cellular pH levels by using the ratiometric pH biosensor NLS-pH-GFP (Moseyko and Feldman, 2001; Fendrych et al., 2014; Lin et al., 2020). Time-lapse imaging showed the onset of nuclear acidification occurred simultaneously with the Ca^2+^ transient in most cells (14/18 cells, Figure 4B and D-E, Figure S3A-B, Movie10). Our results suggest that both Ca^2+^ transient and acidification are early cellular features of dPCD execution in the root cap that coincide with mitochondrial matrix release and precede NE and ER leakage, vacuolar collapse, and PM permeabilization for PI.

**Figure 4:**
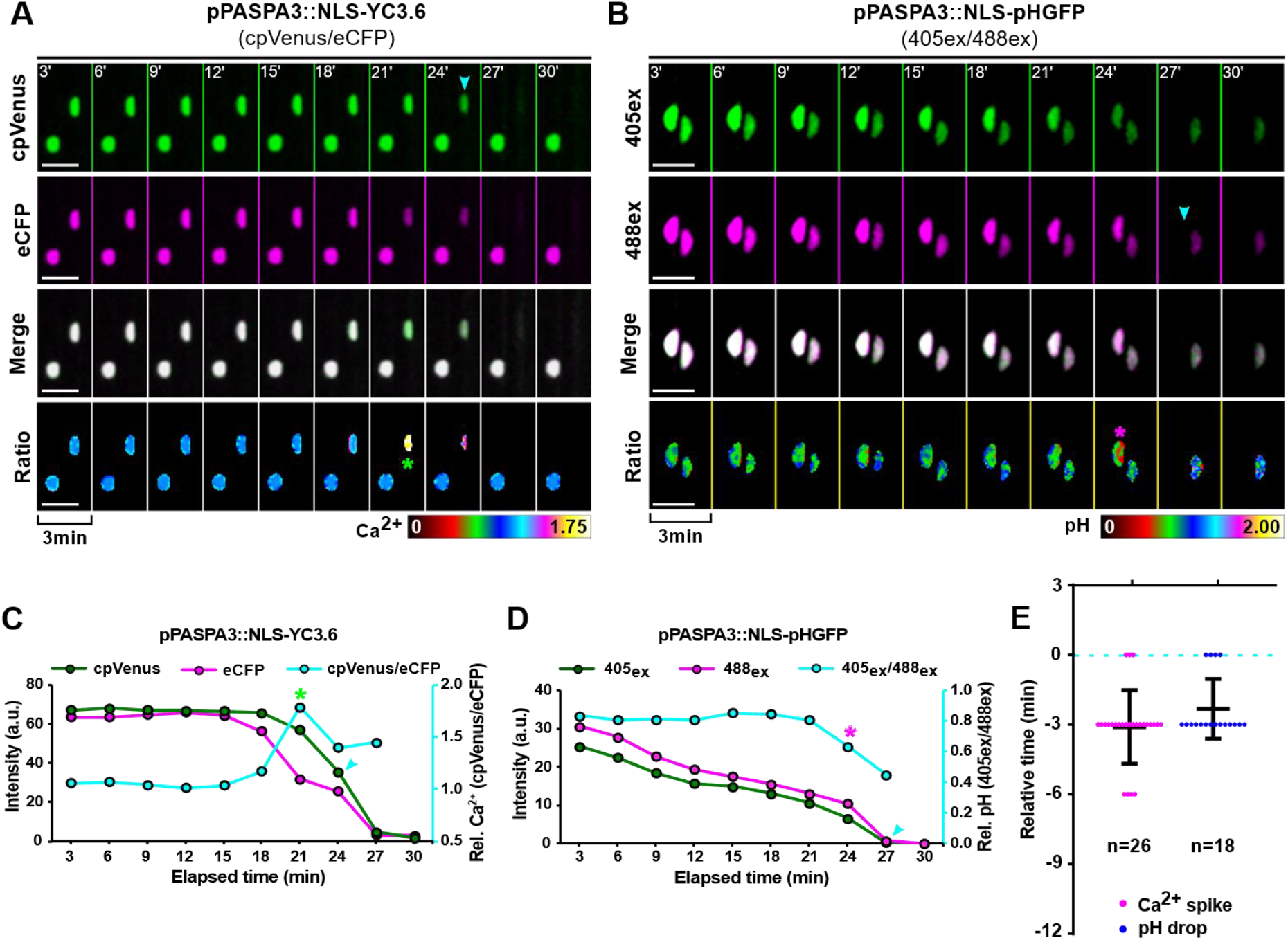
Intracellular calcium transient and acidification occur at the onset of PCD execution. **(A-B)** Confocal Z-projections of time-lapse series in dying LRC cells, cyan arrows indicate NE breakdown. **(A)** False-color ratio images of the VENUS (green) and the CFP (magenta) signals of the calcium sensor *pPASPA3::NLS-YC3.6* indicate a sharp transient increase of Ca^2+^ ions (asterisk) in the nucleoplasm prior to NE breakdown. **(B)** False-color ratio images of the pH sensor *pPASPA3::NLS-pHGFP* show an increase of fluorescence excited at 488 nm (magenta) versus fluorescence excited at 405 nm, indicating an acidification of the nucleoplasm (asterisk) prior to NE breakdown. **(C-D)** Dynamics and ratios of fluorescent intensities and the ratios from Ca^2+^ and pH sensors in a representative cell each. The ratios of fluorescence intensity were obtained after background noise subtraction for each channel and correspond to the relative [Ca^2+^]_nuc_ (cpVENUS/eCFP) or pH_nuc_ (405ex/488ex). The cyan arrow indicates NE breakdown, the green asterisk marks the Ca^2+^ transient and the magenta asterisk marks acidification. **(E)** Quantification of the timing of [Ca^2+^]_nuc_ elevation and onset of pH_nuc_ reduction related to NE breakdown (dashed line); indicated is the mean ± standard deviation, n indicates the number of dying cells analyzed in at least 4 different roots each. Scale bars are 10 μm.

We next investigated whether increased Ca^2+^ concentration and acidification are controlled by the SMB-dependent dPCD pathway in the root cap. Time-lapse imaging of pSMB::NLS-YC3.6 in the *smb-3* mutant indicated that Ca^2+^ elevation prior to cell death still occurs. However, Ca^2+^ elevation occurred earlier than in the wild type, on average 9 minutes before NE leakage became apparent, and was more variable than in the wild type (Figure 5A, C and E, Figure S4A, Movie11). Furthermore, the Ca^2+^ elevation was extended and did not occur in a sharply defined transient as in the wild type (Figure 5A and C). Analyzing dying *smb-3* root cap cells expressing a pSMB::NLS-pHGFP pH biosensor showed a similar pattern for intracellular acidification, indicating that increased Ca^2+^ and H^+^ levels were still associated (Figure 5E). However, intracellular acidification occurred more gradually during *smb-3* cell death (Figure 5 B and D, Figure S4B, Movie12).

**Figure 5:**
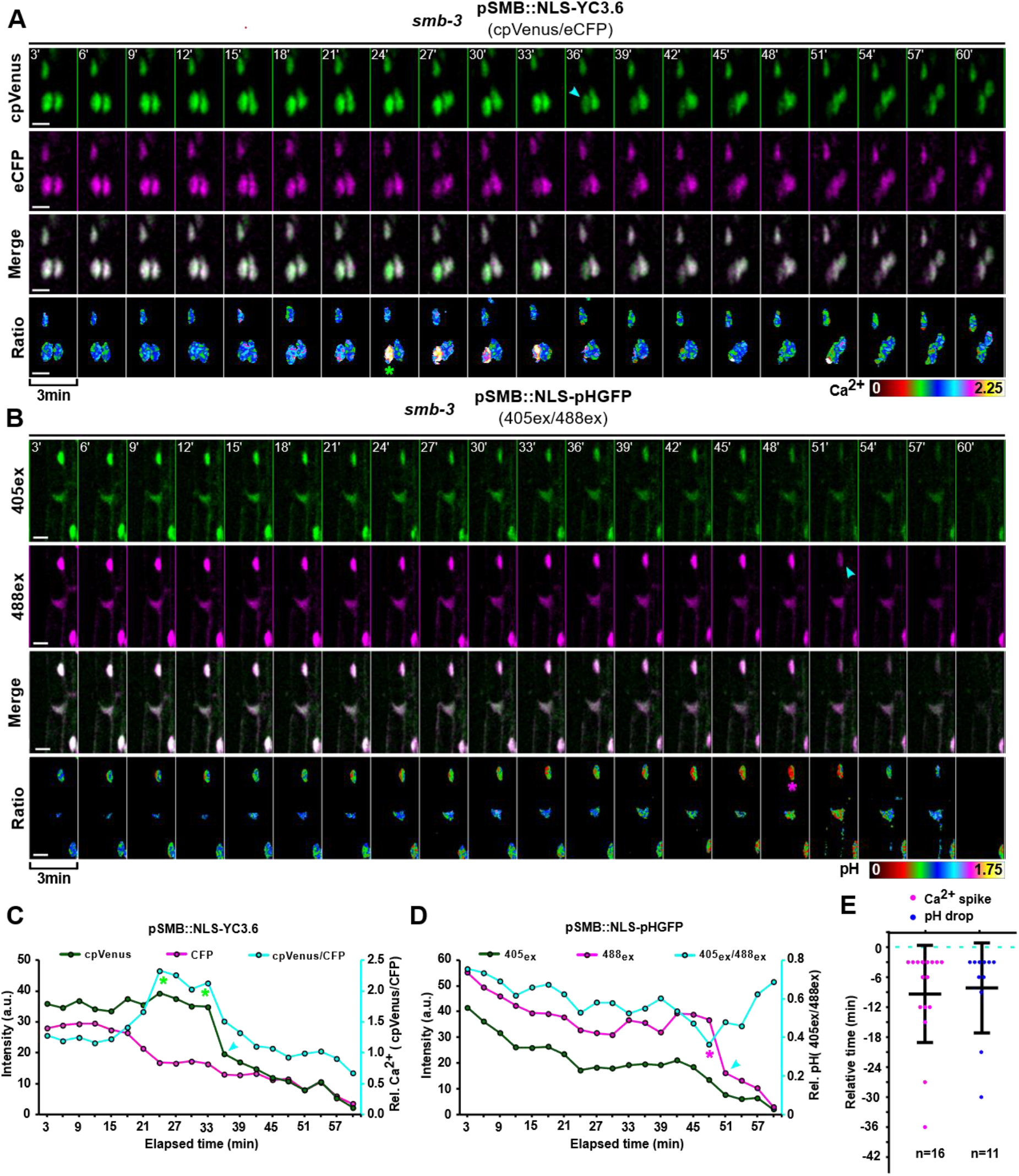
Aberrant patterns of calcium and pH dynamics during *smb-3* cell death. **(A-B)** Confocal Z-projections of time-lapse series in dying *smb-3* LRC cells of 5-day old seedlings. **(A)** False-color ratio images of the calcium sensor *pSMB::NLS-YC3.6* indicate a sustained increase of Ca^2+^ ions (asterisk) in the nucleoplasm prior to NE breakdown (cyan arrow). **(B)** False-color ratio images of the pH sensor *pSMB::NLS-pHGFP* indicate a slow and gradual acidification starting long before NE breakdown (cyan arrow). The pH can reach a minimum (asterisk), but then raises again before NE breakdown. **(C-D)** Dynamics and ratios of fluorescent intensities from Ca^2+^ and pH sensors in a representative *smb-3* cell, each. The ratios of fluorescence intensity were obtained after background noise subtraction for each channel and correspond to the relative [Ca^2+^]_nuc_ (cpVENUS/eCFP) or pH_nuc_ (405ex/488ex). The cyan arrow indicates NE breakdown, the green asterisk marks the start of Ca^2+^ elevation and the magenta asterisk marks the most acidic state of the cell. **(E)** Quantification of the timing of [Ca^2+^]_nuc_ elevation and onset of pH_nuc_ reduction related to NE breakdown (dashed line); indicated is the mean ± standard deviation, n indicates the number of dying cells analyzed in at least 8 different roots each. Scale bars are 10 μm.

These results indicate that Ca^2+^ and H^+^ levels increase during cell death in *smb-3* mutants, but that both events occur in a much slower, aberrant pattern. These findings suggest that in addition to the orchestration of cellular decompartmentalization, SMB targets are involved in controlling the Ca^2+^ signature and intracellular acidification during root cap dPCD.

### Ca^2+^ elevation and acidification are sufficient to trigger cell death in dPCD competent root cap cells

The elevation of intracellular Ca^2+^ and H^+^ could be a mere symptom of membrane modifications at the onset of cell death execution, or be a functional aspect of an efficient dPCD execution. To differentiate between these possibilities, we tested whether pharmacological manipulation of Ca^2+^ and pH levels was sufficient to affect dPCD execution.

Treatment with adenosine 5′-triphosphate (ATP) is widely employed to increase cytoplasmic Ca^2+^ and H^+^ concentrations in *Arabidopsis* seedlings in a dose-dependent manner (Choi et al., 2014; Behera et al., 2018; Waadt et al., 2020). We treated 4-day-old Col-0 seedlings with different doses of ATP in 6-well chambers for 1 hour before mounting them for microscopy. PI staining indicated that a one-hour ATP treatment was sufficient to increase the number of dead root cap cells in a dose-dependent manner (Figure 6A and B). Importantly, cell death was detected mainly in differentiated root cap cells at the edge of the LRC that are preparing for dPCD execution, but not in other cells of the root tip (Figure 6A). In an attempt to differentiate between the effect of increased Ca^2+^ and H^+^ levels, we adjusted the pH during ATP treatment to pH 5.8. This pH-adjusted ATP treatment failed to trigger cell death in root cap cells. However, pH adjustment did not only abolish intracellular acidification, but also interfered with the increase of Ca^2+^ concentrations (Figure 6C-D and Figure S5 E-F). Therefore, we were not able to separate the effects of elevated Ca^2+^ and H^+^ levels on dPCD.

**Figure 6:**
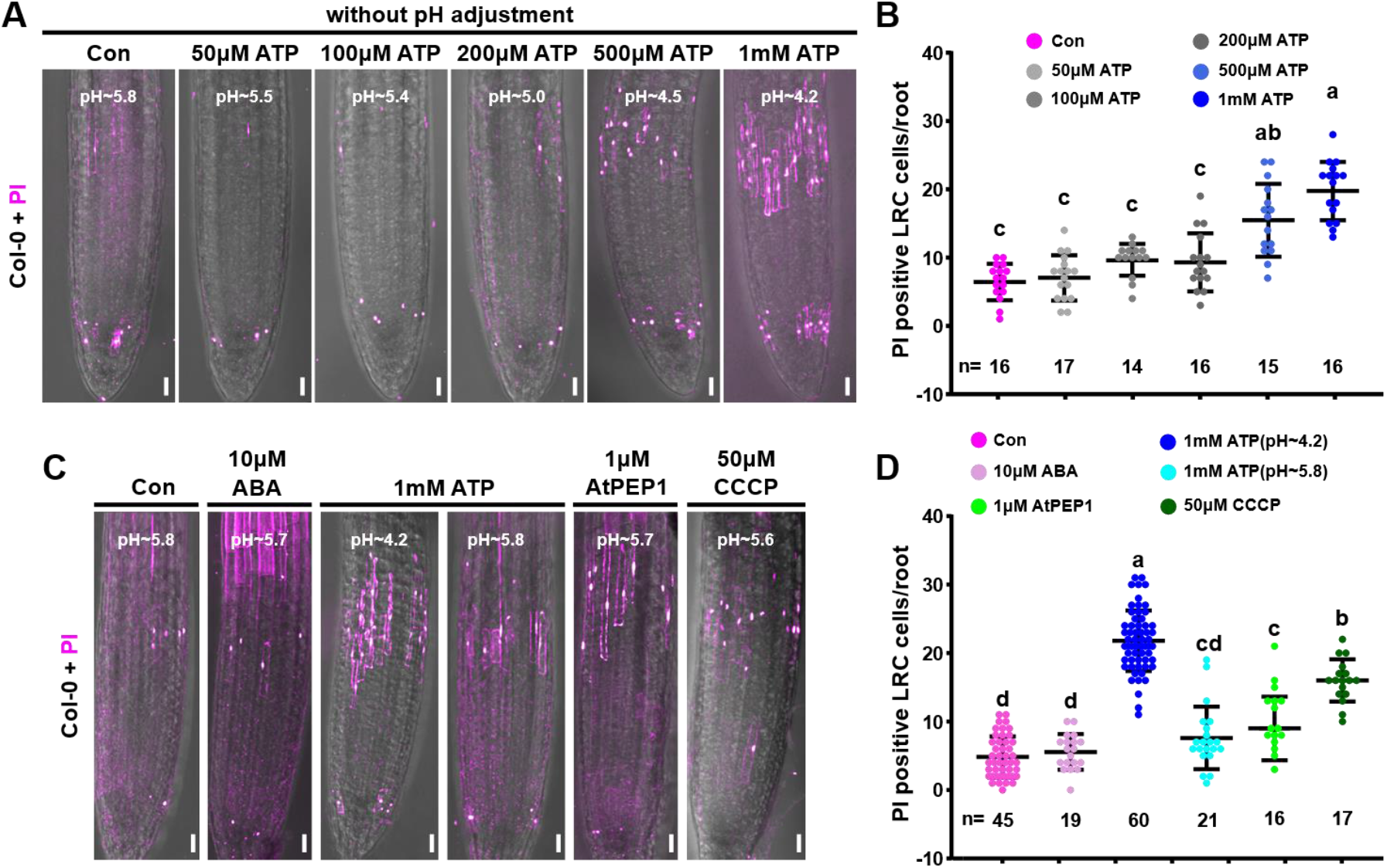
Pharmacological manipulation of pH and intracellular calcium triggers cell death specifically in differentiated root cap cells. **(A)** Confocal Z-projections of PI-stained root tips of 4-day-old Col-0 seedlings after 1h treatments with different concentrations of ATP-Mg. The pH of the ATP solutions is indicated. **(B)** Quantification of PI-positive (dead) cells after a 1h ATP-treatment, indicated is the mean ± standard deviation, n indicates the number seedlings analyzed in each treatment. Letters represent significantly different groups evaluated by one-way anova using Tukey’s multiple comparisons test (P < 0.001). **(C)** Confocal Z-projections of PI-stained root tips of 4-day-old Col-0 seedlings after 1h treatments with different drugs. The pH of the working solutions is indicated. **(D)** Quantification of PI-positive (dead) cells after 1h of treatment, indicated is the mean ± standard deviation, n indicates the number seedlings analyzed in each treatment. Letters represent significantly different groups evaluated by one-way anova using Tukey’s multiple comparisons test (P < 0.001). Scale bars are 20 µm.

To obtain independent pharmacological evidence for the effect of Ca^2+^ and H^+^ manipulation, we used additional molecules that have been described to affect cytoplasmic concentrations of these ions (Waadt et al., 2020). Abscisic acid (ABA) treatment induces cytoplasmic calcium elevation and pH drop in guard cells (Li *et al*., 2021a), but not in Arabidopsis roots (Waadt et al., 2020). In line with these observations, short-term ABA treatment did not affect [Ca^2+^]_nuc_ and pH_nuc_ levels, and failed to trigger dPCD execution in differentiated root cap cells (Figure 6C-D, Figure S5 A-B). By contrast, AtPEP1 and CCCP had been reported to trigger rapid cytosolic acidification and calcium signatures in Arabidopsis roots (Behera et al., 2018; Waadt et al., 2020). Analysis of YC3.6 and pH-GFP upon short-term AtPEP1 and CCCP treatments confirmed this, and both treatments were sufficient to trigger cell death specifically in differentiated root cap cells (Figure 6C-D, Figure S5 G-H). Analysis of YC3.6 and pH-GFP sensors upon CCCP treatment in an incubation chamber system (Krebs and Schumacher, 2013) confirmed the induction of intracellular calcium elevation and acidification in most root cells, and still predominantly differentiated LRC cells reacted with induction of cell death (Supplementary Figure S6 A-B). These observations suggest that differentiated LRC cells are either generally more sensitive to CCCP treatment, or are actively triggering dPCD execution upon sensing elevated Ca^2+^ or H^+^ concentrations.

Together, these results indicate that a short-term increase of intracellular Ca^2+^ and H^+^ concentrations is sufficient to trigger cell death execution specifically in differentiated root cap cells that are prepared to undergo dPCD, while not exerting the same effect in younger LRC or epidermal cells.

Terminal root cap cells in the *smb-3* mutant die in a delayed aberrant cell death that is possibly caused by a physical disruption of cells in the elongation zone (Fendrych et al., 2014). If acquisition of dPCD competency by the SMB gene regulatory network is necessary to react to elevated Ca^2+^ and H^+^ levels with cell death execution, SMB-deficient root cap cells should be insensitive to these signals. To test this hypothesis, we investigated the effect of Ca^2+^ and H^+^ manipulation in *smb-3* mutants. We performed time-course imaging monitoring the dynamics of intracellular [Ca^2+^]_nuc_ and [pH]_nuc_ levels as well as NE leakage as a cell death readout. Firstly, we examined the dynamics of the calcium and pH sensor upon external stimuli in *smb-3* LRC cells. As in the wild type, ABA treatment had no effect on Ca^2+^ and H^+^ levels, and did not lead to an increase of cell death in *smb-3* mutants (Figure 7A-B, Figure S7 A-B). Accordingly, the amount of distalmost root cap cells dying during ABA treatment in *smb-3* mutant is indistinguishable from the wild type (Figure 7A-B). By contrast, short-term CCCP treatment caused a rapid and efficient intracellular calcium elevation and acidification in both wild-type and *smb-3* root cap cells (Figure S5 G-H, Figure S6 C-D). Importantly, however, while CCCP treatment was sufficient to increase cell death specifically in the distal-most root cap cells in the wild type, the distalmost *smb-3* mutant root cap cells showed no increased cell death rates (Figure 7A-B). These results suggest that SMB is necessary to render terminal LRC cells competent to react to elevated Ca^2+^ and H^+^ levels with execution of dPCD.

**Figure 7:**
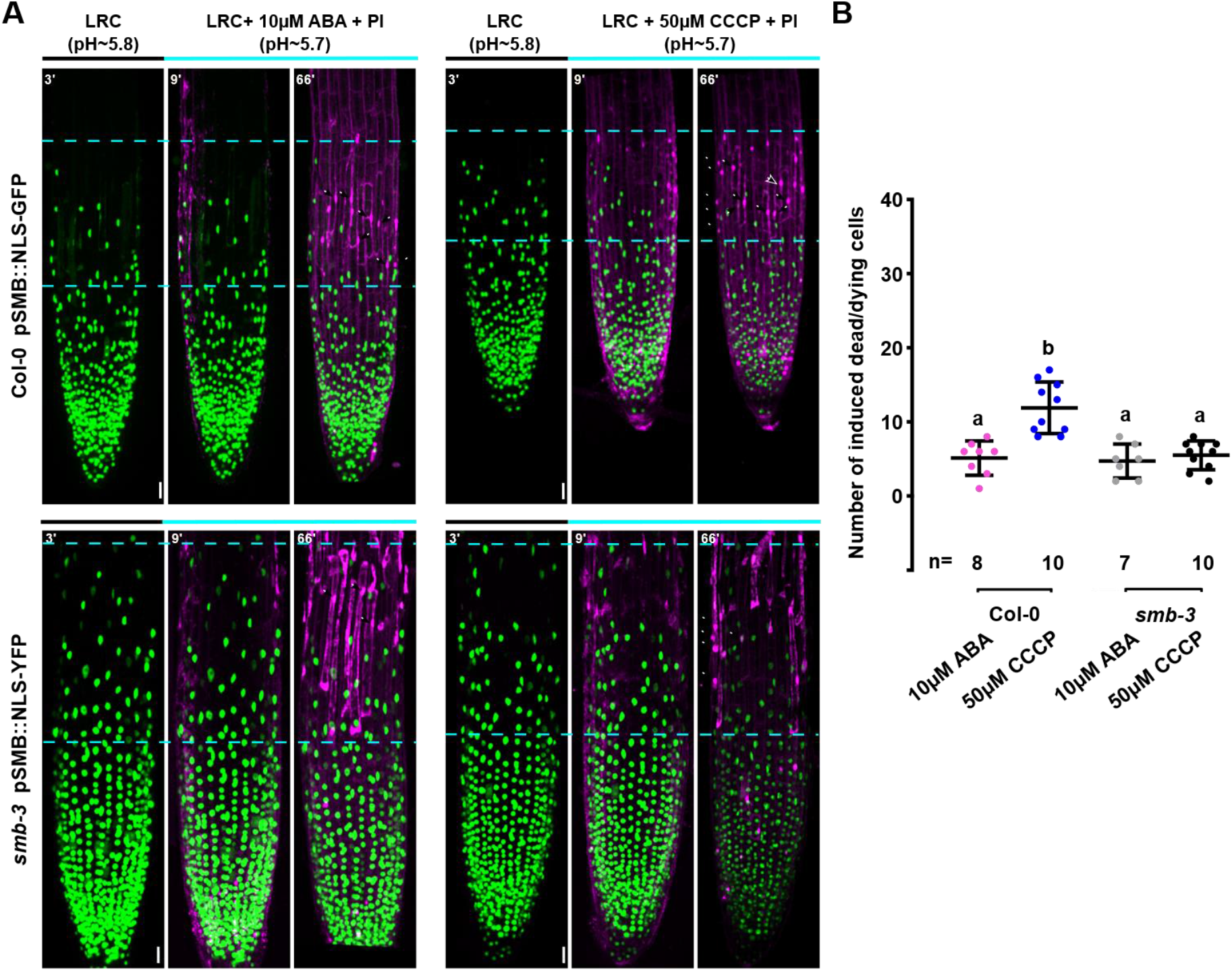
Triggering cell death upon pharmacological manipulation of calcium and pH depends on functional SMB. **(A)** Confocal Z-projections of time-lapse series of LRC cells from 5-day-old Col-0 seedlings expressing *pSMB::NLS-GFP* (upper panels) or *smb-3* seedlings expressing *pSMB::NLS-GFP* (lower panels) stained with PI in response to ABA and CCCP treatment. The distal root cap zone used for quantification of cell death is indicated by dashed lines. Disappearance of NLS-GFP or NLS-YFP signals and appearance of PI signals is used to diagnose cell death after 1h of treatment (white arrows). **(B)** Quantification of dead cells after 1h of treatment, indicated is the mean ± standard deviation, n indicates the number seedlings analyzed in each treatment. Letters represent significantly different groups evaluated by one-way anova using Tukey’s multiple comparisons test (P < 0.01). Scale bars are 20 µm.

## Discussion

The execution of programmed cell death is a remarkably rapid cellular process. In the Arabidopsis root cap it takes only minutes to turn a vital cell into a corpse, and only a few hours to remove the corpse by cell-autonomous clearance (Fendrych et al., 2014). Here we applied multi-factor time-lapse imaging to follow cellular alterations occurring during the dPCD execution in the Arabidopsis root cap. We find that an ordered sequence of selective membrane permeation and ion leakage events takes place during cell death execution. Interestingly, this decompartmentalization process is aberrant in the *smb-3* mutant, suggesting that direct or indirect SMB targets are involved in individual steps of cell death execution (Figure 8).

Cellular decompartmentalization, in particular the collapse of the large central vacuole, has been put forward as a means to drive cell death execution in plants since well over two decades (Fukuda). Vacuolar collapse has been suggested to be “the universal trigger of plant cell death” as it can been observed in several plant cell death contexts (Jones, 2001). Furthermore, vacuolar collapse has been proposed to trigger nuclear degradation based on staining with DNA-intercalating dyes in Zinnia xylogenic cell cultures showing nuclear degradation after vacuolar collapse (Obara et al., 2001). However, we find that vacuolar collapse in the Arabidopsis root cap is a relatively late and temporally variable event that can happen as late as 40 minutes after the NE leakage. By contrast, other membrane modification events leading to NE and ER leakage and the shedding of fluorescent moieties of membrane reporters into the cytosol occur concomitantly on average about 10 minutes before vacuolar collapse.

In the dPCD context, the vacuole has generally been interpreted as a safe storage compartment to accumulate death-executing and corpse-processing hydrolases (Hara-Nishimura and Hatsugai, 2011; Escamez and Tuominen, 2014). However, while vacuolar hydrolases likely contribute to corpse clearance, at least in root cap dPCD they cannot be the exclusive dPCD executioners as vacuolar collapse only occurs after other cellular decompartmentalization events. Several hydrolases that have been implicated in dPCD processes accumulate in the ER lumen prior to dPCD execution, for instance BIFUNCTIONAL NUCELASE1 (BFN1) (Farage-Barhom et al., 2011; Fendrych et al., 2014) or CYSTEINE ENDOPEPTIDASE 1 (CEP1), CEP2, and CEP3 (Hierl et al., 2014; Zhou et al., 2016). Though there is evidence for a direct or indirect transport of ER-resident proteins to the vacuole (Zhang et al., 2014; Toyooka et al., 2023), ER leakage releases high quantities of such hydrolytic enzymes into the cytosol prior to vacuolar collapse. Thus, in combination with intracellular acidification, ER leakage may activate several hydrolytic enzymes that are known to be pH dependent, for instance CEP1 (Hierl et al., 2014). In this scenario, ER leakage would start a proteolytic cascade that triggers an irreversible cell death execution process leading to vacuolar collapse and subsequent corpse clearance. It will be an intriguing challenge for future research to test if ER-resident proteins are responsible for vacuolar collapse.

Intriguingly, however, ER and NE leakage is preceded by the release of mitochondrial matrix proteins into the cytosol. In animal apoptosis, the permeabilization of the mitochondrial outer membrane is a key step in the intrinsic pathway, releasing cytochrome c and pro-apoptotic factors from the mitochondrial intermembrane space that trigger a caspase-cascade leading to apoptotic cell death (Tait and Green, 2013). More recently, BAX/BAK macropore formation in the mitochondrial outer membrane have been shown to lead to rupture of the mitochondrial inner membrane and release of matrix factors, including mitochondrial DNA, into the cytosol (McArthur et al., 2018; Riley et al., 2018). Our data suggest that a similar mechanism might operate in plant dPCD, despite the absence of BAX/BAK orthologs. Mitochondrial involvement in plant cell death has been controversially discussed (Minina et al., 2021); cytochrome c release has been reported in several PCD contexts (Balk et al., 1999; Balk and Leaver, 2001; Thomas and Franklin-Tong, 2004), though cytochrome c as such has been found insufficient for PCD execution (Yu et al., 2002).

At the onset of cellular decompartmentalization, we observed characteristic elevations of intracellular Ca^2+^ and H^+^ levels. Changes in calcium and pH levels have been observed in a wide variety of plant PCD processes and been discussed as potential death triggers (Jones, 2001; Ren et al., 2021). For instance, cytoplasmic Ca^2+^ influx in Zinnia xylogenic cell cultures has been shown to promote PCD. Treatment with EGTA (chelating extracellular Ca^2+^) or lanthanum (inhibiting Ca^2+^ influx) caused a moderate decrease of cell death induced by the toxin mastoparan. Conversely, ionophore treatment increased cell death rates in a Ca^2+^ dependent manner (Groover and Jones, 1999). Ca^2+^ influx has also been reported from self-incompatibility PCD occurring in poppy (*Papaver rhoeas*) (Wu et al., 2011; Wang et al., 2020). We show that the pattern and efficiency of Ca^2+^ and H^+^ increase depends on the presence of the transcription factor SMB. Furthermore, pharmacologically induced increase of Ca^2+^ and H^+^ levels is sufficient to trigger cell death execution in a SMB-dependent fashion specifically in terminally differentiated root cap cells. Intracellular calcium elevation and acidification are closely associated (Behera et al., 2018; Waadt et al., 2020), and potentially both events could activate processes that lead to selective membrane permeation events. However, since changes in Ca^2+^ and H^+^ levels occur simultaneously with early decompartmentalization events, it remains a challenge to discern between causes and consequences in this context.

There has been a controversial discussion on the presence or absence of apoptosis-like pathways in plant PCD (Minina et al., 2021). Our results demonstrate the maintenance of PM integrity during post-mortem corpse clearance, and the formation of extracellular vesicles (EVs) that are blebbing out of root cap cell corpses. While the formation and morphology of these EVs and apoptotic bodies are probably fundamentally different (Battistelli and Falcieri, 2020; Santavanond et al., 2021), they might serve a similar function: To prevent the leakage of cellular components to the extracellular space. In apoptosis this is part of a sophisticated strategy to prevent inflammation by avoiding the release of pro-inflammatory signals and generating signals that suppress inflammation (Yamaguchi et al., 2014). In plants, which do not produce inflammatory responses, the maintenance of PM impermeability for proteins might serve a different function, preventing the release of hydrolytic enzymes into the apoplast. In the acidic pH of the extracellular space, such enzymes could damage proteins and other molecules present in the cell wall and the PM of neighboring epidermal cells. While testing this hypothesis remains challenging, the maintenance of PM integrity remains an intriguing possible parallel between specific animal and plant cell death routines.

Summing up, it is tempting to speculate that the cellular mechanism that facilitates cell death execution in the Arabidopsis root cap consists of a series of genetically controlled selective and successive decompartmentalization events. The root cap is arguably more accessible for live-cell imaging and pharmacological manipulation than most other tissues or cell types undergoing dPCD in plants, but it will be an interesting task to investigate if the sequence of events we observed in the root cap is reproducible in other dPCD processes. Furthermore, figuring out how the individual cellular membranes are differentially permeabilized during dPCD execution remains a future challenge. Ultrastructural analyses, for instance guided by correlative light and electron microscopy, will be necessary, but it will remain challenging to visualize the rapid cellular changes of dPCD execution on the ultrastructural level. Last, but not least, to identify the proteins and the mechanisms by which they achieve controlled membrane permeabilization will be crucial to understand dPCD execution in plants.

## Materials and Methods

### Molecular cloning

Expression constructs used in this study were summarized in Supplementary Table S1. Expression vectors included in this study that have been previously described elsewhere include pPASPA3::H2A-GFP (Fendrych et al., 2014), pPASPA3::NLS-TdTOMATO (Xuan et al., 2016), pPASPA3::ToIM (Fendrych et al., 2014), pSMB::NLS-GFP-GUS (Xuan et al., 2016), pSMB::NLS-YFP (Kamiya et al., 2016), pUBQ10::GFP-VAMP711 (Feng et al., 2017), and 35S::GFP-GPI (Martiniere et al., 2012). The pUBQ10::3xmCHERRY-SYP122 was cloned based on the pUBQ10 promoter sequence in the GreenGate collection pGGA006 (Lampropoulos et al., 2013) via KpnI upstream of the 3xmCHERRY-SYP122 reporter gene (Andersen et al., 2018) in pGreen179.

Gateway entry vectors were generated by amplifying the desired DNA fragment with appropriate attB recombination sites, and subsequently inserting into a proper donor vector using BP Clonase enzyme (ThermoFisher). The details of primers used in this study were listed in Supplementary Table S2. Gateway expression vectors were generated by recombining entry vectors into specified destination vectors using LR Clonase II plus enzyme (ThermoFisher).

Double gateway LR reactions (Invitrogen) were employed to obtain the expression vectors pPASPA3::NLS-pHGFP and pPASPA3::NLS-YC3.6. Primers PR12/13 were used to amplify pHGFP, including an N-term NLS, using pUBQ10::pHGFP (Fendrych et al., 2014) as template. attB1 and attB2 sites were added by PCR using PR14/15 with the above PCR product as template. The resulting PCR fragment was cloned into pDONR221 to obtain entry clone of pEN-L1-NLS-pHGFP-L2. Primers PR16/17 were used to amplify NLS-YC3.6 using pUBQ10::NLS-YC3.6 (Krebs et al., 2012) as template. The resulting DNA fragment was cloned into pDONR221 to obtain entry clone of pEN-L1-NLS-YC3.6-L2. pEN-L4-pPASPA3-R1 (Fendrych et al., 2014), pEN-L1-NLS-pHGFP-L2 and pEN-L1-NLS-YC3.6-L2 were recombined into pB7m24GW (Karimi et al., 2002) respectively.

The COX4-mts fragment was first originally described elsewhere (Nelson et al., 2007). To obtain different promoter driven COX4-mts-mTFP1 fusions, primers PR5/6 were used to amplify mTFP1 fragment, insert the COX4 mitochondrial targeting sequence (mts) before mTFP1, and add attB1 and attB2 sites using pEN-R2-mTFP1 2xFLAG-L3 as template. The resultant DNA fragment was cloned into pDONR221 to obtain entry clone of pEN-L1-COX4-mts-mTFP1-L2. Primers PR18/19 were used to amplify 1,169bp promoter fragment before the start codon of TATD (At3g03500) from gDNA, and the resulting PCR fragment was cloned into pDONRP4-P1R to obtain pEN-L4-pTATD-R1. Entry clones of pEN-L4-pH3.3-R1 (Ingouff et al., 2017) or pEN-L4-pTATD-R1, and pEN-L1-cox4-mts-mTFP1-L2 were recombined into pB7m24GW (Karimi et al., 2002) or pB7m24GW-FAST-Green (https://gatewayvectors.vib.be) to obtain expression vectors respectively mTFP1.

The CPK17 fragment was originally described elsewhere (Benetka et al., 2008). Primers PR7/8 were used to amplify CPK17 fragment and add attB1 and attB2 sites, using template described before (Mehlmer et al., 2012). The resultant DNA fragment was cloned into pDONR207 to obtain pEN-L1-CPK17 fragment-L2. Primers PR20/21 were used to amplify mTFP1 (with 2xFLAG tag) sequence and add attB2 and attB3 sites, using template from Pasin and colleagues (Pasin et al., 2014). This PCR product was cloned into pDONRP2-P3R to obtain pEN-R2-mTFP1-2xFLAG-L3. Then the entry clones of pEN-L4-pH3.3-R1 (Ingouff et al., 2017) or pEN-L4-pPASPA3-R1 (Fendrych et al., 2014), pEN-L1-CPK17 fragment-L2, and pEN-R2-mTFP1-2xFLAG-L3 were recombined into pB7m34GW (Karimi et al., 2007) or pB7m24GW,3 (Karimi et al., 2002) to obtain expression vectors respectively.

Primers PR9/10 were used to amplify mTFP1 fragment, insert C-term KDEL and N-term SP sequences, and add attB2 site using pEN-R2-mTFP1 2xFLAG-L3 as template. This fragment was further amplified using PR9/11 to add attB1 site, and cloned into pDONR207 to obtain pEN-L1-SP-mTFP1-KDEL-L2. The entry clones of pEN-L4-pH3.3-R1 (Ingouff et al., 2017) or pEN-L4-pTATD-R1, and pEN-L1-SP-mTFP1-KDEL-L2 were recombined into pB7m24GW,3 (Karimi et al., 2002) to obtain expression vectors respectively.

Primers PR22/23 were used to amplify 2.5kb promoter fragment before the start codon of SCAMP5 (At1g32050) between attB4 and attB1 sites. The resulting PCR fragment was cloned into pDONRP4-P1R to obtain pEN-L4-pSCAMP5-R1. Primers PR24/25 were used to amplify genomic SCAMP5 (without stop codon) between attB1 and attB2 sites. The resulting PCR fragment was cloned into pDONR221 to obtain pEN-L1-pSCAMP5-L2. Entry clones of pEN-L4-pSCAMP5-R1, pEN-L1-SCAMP5-L2, and pEN-R2-eGFP-L3 were recombined into pB7m34GW (Karimi et al., 2007) to obtain the expression vector pSCAMP5::SCAMP5-eGFP.

pEN-L4-pH3.3-R1 (Ingouff et al., 2017) and pEN-L1-NLS-TdTOMATO-L2 (Xuan et al., 2016) were recombined into pB7m24GW-FAST-Green (https://gatewayvectors.vib.be) to obtain the expression vector pH3.3::NLS-TdTOMATO.

Golden Gate entry modules pGG-A-pSMB-B, pGG-B-Linker-C were previously reported (Decaestecker et al., 2019). pGG-D-Linker-E, pGG-E-tHSP18.2M-F, pGG-F-linkerII-G were collected from PSB plasmids stocks. Primers PR1/2 were used to amplify NLS-pHGFP using pEN-L1-NLS-PHGFP-L2 (described above) as a template, and the subsequent fragment was inserted into BsaI-digested GreenGate entry vector pGGC000 (Lampropoulos et al., 2013) via Gibson assembly to generate pGG-C-NLS-pHGFP-D. Primers PR3/PR4 were used to amplify NLS-YC3.6 using pEN-L1-NLS-YC3.6-L2 (described above) as a template, and the subsequent fragment was inserted into BsaI-digested GreenGate entry vector pGGC000 (Lampropoulos et al., 2013) via Gibson assembly to generate pGG-C-NLS-YC3.6-D. Entry modules were assembled in pFASTRK-AG via Golden Gate reactions, resulting in the expression vectors pSMB::NLS-pHGFP and pSMB::NLS-YC3.6.

PCRs for cloning were performed using Phusion high-fidelity DNA polymerase (ThermoFisher). All entry and expression vectors were checked by Sanger sequencing, using Eurofins Scientific Mix2Seq or TubSeq services. Information regarding plasmids and primers can be found in Supplemantary Tables S1 and S2, respectively.

### Arabidopsis transgenic plants and growth conditions

All plant materials used in this study are in Col_0 ecotype background. Information on plants materials is listed in Supplementary Table 3. To generate the single-color reporter lines or marker lines, Col_0 *or smb-3* plants were transformed with the expression constructs respectively by floral dipping. Primary transformants were selected with antibiotics selection or FASTR selection. The FASTR-positive T1 transgenic seeds were selected under a Leica M165FC fluorescence stereomicroscope using the DSR fluorescence filter (excitation: 510-560nm; emission 590-650nm). To generate the dual-color marker lines, the T2 or T3 generation of Col_0 or *smb-3* plants expressing single fluorescent markers were selected on SP8x confocal with nice fluorescent signals and used for crossing respectively. The F1 or F2 generation of dual-color marker lines in Col_0 or *smb-3* background were used for imaging.

Seeds were sterilized by chlorine gas sterilization and sown on LRC medium (1/2 MS [Duchefa Biochemie], 0.1 g/L MES, pH 5.8 [KOH], and 0.8% plant agar [Neogen]) following a 3-day vernalization period at 4°C. Then seeds were moved to growth chamber for vertical growth with continuous light emitted by white fluorescent lamps (intensity of 120 μmol m^-2^ s^-1^) at 22°C for 4 or 5 days.

### Live-Cell Imaging, staining and pharmacological treatment

A Leica SP8X confocal microscope equipped with a white light laser (WLL) was used for all confocal imaging, except the imaging of pPASPA3::ToIM (Figure 2E), which that was performed on a Zeiss 900 vertical microscope. Confocal imaging on SP8X was acquired with a 40x (HC PL APO CS2, NA=1.10) or a 25x (HC FLUOTRA, NA=0.95) water-immersion corrected objective. Confocal imaging on the vertically mounted Zeiss 900 was performed with a 20x dry objective.

For the reporter lines or marker lines, the GFP, mTFP1, YFP, mCHERRY and TdTOMATO fluorescent proteins were excited with 488nm, 470nm, 514nm and 561nm WLL laser lines respectively. And the fluorescence emissions were collected as below: GFP (500-530 nm), mTFP1 (480-530 nm), YFP (520-550 nm), mCHERRY and TdTOMATO (580-650 nm). For the radiometric biosensors, the pH sensor pHGFP were excited with 405 nm and 488 nm, and fluorescence emissions were collected between 500 and 540 nm. The calcium sensor YC3.6 was excited with 405 nm, and the fluorescence emissions were collected with 460-500 nm (for eCFP) and 520-545 nm (for cpVenus). When seedlings were performed with PI (10 μg/mL, Sigma-Aldrich) or FM4-64 (4μM, Invitrogen) staining, dyes were excited with 561 nm and emissions were collected between 600 and 700 nm. Confocal images on SP8X were acquired with Hybrid detectors (HyDTM) using a time-gated window between 0.3ns-6.0ns (for fluorescent proteins) and in a line sequential mode. For pPASPA3::ToIM, images were acquired as previously described (Fendrych et al., 2014) on a vertically mounted Zeiss 900 using a 20x objective.

Time-course confocal imaging (xyzt) was performed to monitor the dynamic behaviors of reporters or biosensors. To obtain confocal imaging for Figure 1-5 and Figure S1-S2, seedlings expressing single or dual-color reporter lines, NLS-pHGFP and NLS-YC3.6 biosensors in Col_0 (4-day-old) and *smb-3* (5-day-old) background were transferred to a Lab-Tek chamber and covered with an agar slab (LRC) supplemented with or without 10 μg/mL PI. To maintain the humidity, before mounting the seedlings, the Lab-Tek chamber meanwhile was supplemented with appropriate amount (100 ∼200 μL) of LRC liquid medium (1/2 MS [Duchefa Biochemie], 0.1 g/L MES, pH 5.8 [KOH]) with or without 10 μg/mL PI.

To explore the efficiency of external stimuli upon induction of cell death in LRC cells (Figure 6), 4-day old Col_0 seedlings were incubated with LRC liquid medium supplemented with indicated concentrations of compounds (Adenosine 5′-triphosphate magnesium salt, ATP-Mg [VWR]; Abscisic acid, ABA [Sigma-Aldrich]; Carbonyl cyanide 3-chlorophenylhydrazone, CCCP [Sigma-Aldrich]; AtPEP1 [kindly provided by Prof. Eugenia Russinova] in 6-well plates for 1h, following with shortly PI-staining (10 μg/mL PI in the incubated medium) for 2mins. The treated seedlings were transferred into Lab-Tek chamber and covered with an agar slab (LRC), by supplementing adequate incubated medium. Z-projections images were acquired SP8x confocal, and the whole imaging process of treated seedlings was accomplished within 20 mins. Before the incubation of seedlings, the pH of incubated medium was measured. For the pH adjustment of ATP treatment, KOH was used to adapt pH to 5.8. PI-stained positive LRC cells of the whole roots were counted.

A microscope incubation chamber system (Krebs and Schumacher, 2013) was employed to monitor the dynamics of relative calcium and pH level upon different external stimuli application (Figure S5 and Figure S6), as well as evaluate the induction of cell death in Col_0 and *smb-3* seedlings. To study the dynamic behaviors of [Ca^2+^] _nuc_ and pH _nuc_ upon external stimuli treatment, Col_0 seedlings (4-day-old) or *smb-3* seedlings (5-day-old) expressing [Ca^2+^] _nuc_ and pH _nuc_ biosensors were placed in the dedicated chamber and covered with fiberglass soaked in imaging solution (200 μL LRC liquid medium). During the time-course imaging, treatment was performed by quickly supplementing the imaging solution with same volume of LRC liquid medium supplemented with 2x concentrations of indicated compounds and PI. The PI staining was used to indicate the application of external stimuli. Similarly, comparisons between ABA- and CCCP-induced cell death in Col_0 and *smb-3* seedlings were also performed with the microscope incubation chamber system (Figure 7). The Col_0 seedlings and *smb-3* seedlings grown on LRC plates for 5days were used for experiments. Time-course confocal imaging experiments were performed to monitor the induced dying/dead cells upon ABA and CCCP application in Col_0 and *smb-3* seedlings. Treatment was performed by quickly supplementing the imaging solution with same volume of LRC liquid medium supplemented with 2x concentrations of indicated compounds and PI. The PI staining was used to indicate the application of external stimuli as well as stained dead LRC cells. Disappearance of NLS-GFP or NLS-YFP signals and appearance of PI signals was used to diagnose cell death. Only the distalmost LRC cells of Col-0 and *smb-3* expressing strong NLS-GFP or NLS-YFP signal upon treatment were counted.

### Image analysis and figure preparation

Image processing and analyses were conducted using LAS X and Fiji (Schindelin et al., 2012). To compensate for the root growth, the time-lapse images were registered using the “rigid body” options of the MultiStackReg (https://github.com/miura/MultiStackRegistration). The processed images from Image J were cropped and assembled for figures in Inkscape (http://inkscape.org).

For the analysis of relative calcium and pH (NLS-YC3.6 and NLS-pHGFP images) level during cell death, Z-projections were generated from time-course confocal images using the “Maximum projection” of LAS X and registered using Fiji. Analysis of NLS-YC3.6 and NLS-pHGFP were described in details before (Doccula et al., 2018). Briefly, intensities of cpVenus and eCFP (for YC3.6), or 405_em_ and 488_em_ (for pHGFP) emissions of the analyzed regions of interest (ROIs) corresponding to the entire nucleus of the intrested cell and used for the ratio (R) calculation (cpVenus/eCFP; 405_ex_/488_ex_). Before ratio calculation, background noises for each channel of fluorescence signals were subtracted. The background noises were determined by the background signal of each channel and corresponded to a ROI chosen next to the nucleus of interest. Fluorescence intensities and background noises of each channel during cell death were documented and plotted in excel. The sudden drop of nucleus fluorescence intensities of cpVenus and 488_ex_ were used to determine as the timing of NE breakdown. Ratios dynamics of cpVenus/eCFP and 405_ex_/488_ex_ were obtained as below and reflect the dynamics of relative Ca^2+^ and pH level during cell death respectively (Figure 4 and 5, Figure S3 and S4).

Rel. Ca^2+^ (cpVenus/eCFP) = (cpVenus - cpVenus background)/ (eCFP-eCFP background)

Rel. pH (405_ex_/488_ex_) = (405_ex_ - 405_ex_ background)/ (488_ex_ - 488_ex_ background)

For the ratio images analysis of NLS-YC3.6 and NLS-pHGFP images (Figure 4 and 5, Figure S5-6), fluorescence intensities of the selected ROIs were extracted using Fiji for both the eCFP and cpVenus channels, or for both of 405_ex_ and 488_ex_ channels. The “Otsu” method of threshold establishment was used to generate masks of the images. The extracted eCFP and cpVenus channels, or 405_ex_ and 488_ex_ channels were normalized to the same mask images respectively by using the “Image Calculator” tool respectively. The ratiometric images were obtained as below by reusing the “Image Calculator” tool respectively.

Ratio images of YC3.6 = (extracted cpVenus/mask)/ (extracted eCFP/mask)

Ratio images of pHGFP = (extracted 405_ex_ /mask)/ (extracted 488_ex_ /mask)

### Statistical analysis

Statistical analysis was performed using GraphPad Prism 9.0.0 for Windows and statistical details of experiments were indicated in the corresponding figure legends.

## Author contributions

J.W. performed the research and analyzed data; N.B., and R.A.B. designed the study, and together with R.H. and H.V. generated marker lines; D.V.D. designed and supervised experiments, E.M. contributed to live-cell imaging and experimental setup; Q.J. generated the SCAMP5-GFP line; Z.L. contributed to the construction of vector and transgenic lines; M.K.N. supervised and designed the study and wrote the manuscript together with J.W. and H.V.

## Acknowledgements

We would like to thank all members of the PCD lab at the VIB-UGent Center for Plant Systems Biology for discussions and critical comments. We gratefully acknowledge funding from the China Scholarship Council (CSC grant No. 201906760018 to Q.J.), the European Research Council (ERC StG PROCELLDEATH 639234 and CoG EXECUT.ER 864952 to M.K.N.), the Research Foundation – Flanders (FWO project No. G041118N, and FWO postdoctoral fellowship No. 12I7420N to Z.L.), and the University of Ghent Bijzonder Onderzoeksfonds (BOF postdoctoral fellowship No. 01P06118 to Q.F.).

## Movie Legends

Movie1: Time-course confocal imaging indicates NE breakdown as reference time point during dPCD execution. Time-course confocal imaging of a root tip from a 4-day-old Col-0 seedling expressing the nuclear reporter *pSMB::NLS-GFP* stained with PI. The NLS-GFP signal dissipates in the distalmost LRC cell (arrowhead), indicating the rapid occurrence of NE breakdown, followed by PI entry into the cell (magenta signal). A maximal Z-projection of 3 slices is shown. Root growth was compensated for by registration. Related to Figure 1.

Movie2: Time-course confocal imaging indicates NE breakdown occurs concurrently with ER leakage during dPCD. Time-course confocal imaging of a root tip from a 4-day-old Col-0 seedling expressing *pPASPA3::NLS-TdTOMATO* and *pTATD::SP-mTFP1-KDEL* dual-color markers. In the dying cells of the distalmostdistalmost LRC cells, NE breakdown is accompanied by ER leakage into the cytosol, preceding vacuolar collapse indicated by even distribution of the fluorescent signal in the entire cell volume. A maximal Z-projection is shown. Root growth was compensated for by registration. Related to Figure 1.

Movie3: Time-course confocal imaging indicates mitochondrial matrix leakage prior to NE breakdown during dPCD. Time-course confocal imaging of a root tip from a 4-day-old Col-0 seedling expressing *pPASPA3::NLS-TdTOMATO* and *pTATD::COX4-mTFP1* dual-color markers. In the dying cells of the distalmost LRC cells, mitochondria leakage, indicated by abrupt disappearance of mitochondrial foci, occurs prior to NE breakdown. A maximal Z-projection is shown. Root growth was compensated for by registration. Related to Figure 1.

Movie4: Time-course confocal imaging indicates NE breakdown occurs concurrently with PM endodomain shedding during dPCD. Time-course confocal imaging of a root tip from a 4-day-old Col-0 seedling expressing *pPASPA3::NLS-TdTOMATO and pPASPA::CPK17-mTFP1* dual-color markers. During dPCD in the distalmost LRC cells, NE breakdown occurs simultaneously with PM endodomain shedding (indicated by strong cytoplasmic signal), and prior to vacuolar collapse (indicated by an even distribution of the fluorescent protein in the entire cell volume). A maximal Z-projection is shown. Root growth was compensated for by registration. Related to Figure 2.

Movie5: Time-course confocal imaging indicates generation of extracellular vesicles during dPCD. Time-course confocal imaging of a root tip from a 4-day-old Col-0 seedling expressing *pPASPA3::ToIM*. During dPCD occurring in the distalmost LRC cells, extracellular vesicles can be seen blebbing out of degrading cell corpses on the root surface. Note that living cells show a strong cytoplasmic GFP signal (green) and degrading cell corpses show an evenly distributed RFP signal (magenta). A maximal Z-projection is shown. Root growth was compensated for by registration. Related to Figure 2.

Movie6: Time-course confocal imaging indicates gradual dissipation of nuclear localized fluorescent proteins during cell death in the *smb-3* mutant. Time-course confocal imaging of a root tip from a 5-day-old *smb-3* seedling expressing the nuclear reporter *pSMB::NLS-GFP* stained with PI. In the dying LRC cell (nucleus indicated by arrowhead), the NLS-GFP signal disappears more gradually than in the wild type, followed by PI entry. A maximal Z-projection is shown. Root growth was compensated for by registration. Related to Figure 3.

Movie7: Time-course confocal imaging indicates delayed mitochondrial matrix release during cell death in the *smb-3* mutant. Time-course confocal imaging of a root tip from a 5-day-old *smb-3* seedling expressing *pH3.3::NLS-TdTOMATO* and *pH3.3::COX4-mTFP1* dual-color markers. In the dying LRC cells, mitochondrial foci remain visible after NE breakdown and vacuolar collapse. A maximal Z-projection is shown. Root growth was compensated for by registration. Related to Figure 3.

Movie8: Time-course confocal imaging indicates the aberrant PM endodomain shedding during cell death in the *smb-3* mutant. Time-course confocal imaging of a root tip from a 5-day-old *smb-3* seedling expressing *pH3.3::NLS-TdTOMATO* and *pH3.3::CPK17-mTFP1* dual-color markers. In the dying LRC cells, cytosolic solubilization of CPK17-mTFP1 is only completed after NE breakdown in *smb-3* mutants. A maximal Z-projection is shown. Root growth was compensated for by registration. Related to Figure 3.

Movie9: Time-course confocal imaging indicates the intracellular calcium transient during dPCD. Time-course confocal images (left) and false-color ratio images (right) from a 4-day-old Col-0 seedling expressing the calcium sensor *pPASPA3::NLS-YC3.6*. In the dying distal LRC cells, a sharp transient increase of Ca^2+^ ions in the nucleoplasm occurs prior to NE breakdown. A maximal Z-projection of 3 slices is shown. Root growth was compensated for by registration. Related to Figure 4.

Movie10: Time-course confocal imaging indicates the intracellular acidification during dPCD. Time-course confocal images (left) and false-color ratio images (right) from a 4-day-old Col-0 seedling expressing the pH sensor *pPASPA3::NLS-pHGFP*. In the dying distal LRC cells, acidification of the nucleoplasm occurs prior to NE breakdown. A maximal Z-projection of 3 slices is shown. Root growth was compensated for by registration. Related to Figure 4.

Movie11: Time-course confocal imaging indicates the aberrant patterns of intracellular calcium transient during *smb-3* cell death. Time-course confocal images (left) and false-color ratio images (right) from a 5-day-old *smb-3* seedling expressing a calcium sensor *pSMB::NLS-YC3.6*. In the dying distalmost LRC cells, an increase of Ca^2+^ ions in the nucleoplasm is followed by a gradual occurrence of NE breakdown. A maximal Z-projection from 3 slices is shown. Root growth was compensated for by registration. Related to Figure 5.

Movie12: Time-course confocal imaging indicates the aberrant patterns of intracellular calcium transient during *smb-3* cell death. Time-course confocal images (left) and false-color ratio images (right) from a 5-day-old *smb-3* seedling expressing a pH sensor *pSMB::NLS-pHGFP*. In the dying distalmost LRC cells, a slow and gradual acidification occur before NE breakdown (cyan arrow). A maximal Z-projection of 3 slices is shown. Root growth was compensated for by registration. Related to Figure 5.

---

**Supplemental Figure S1:**
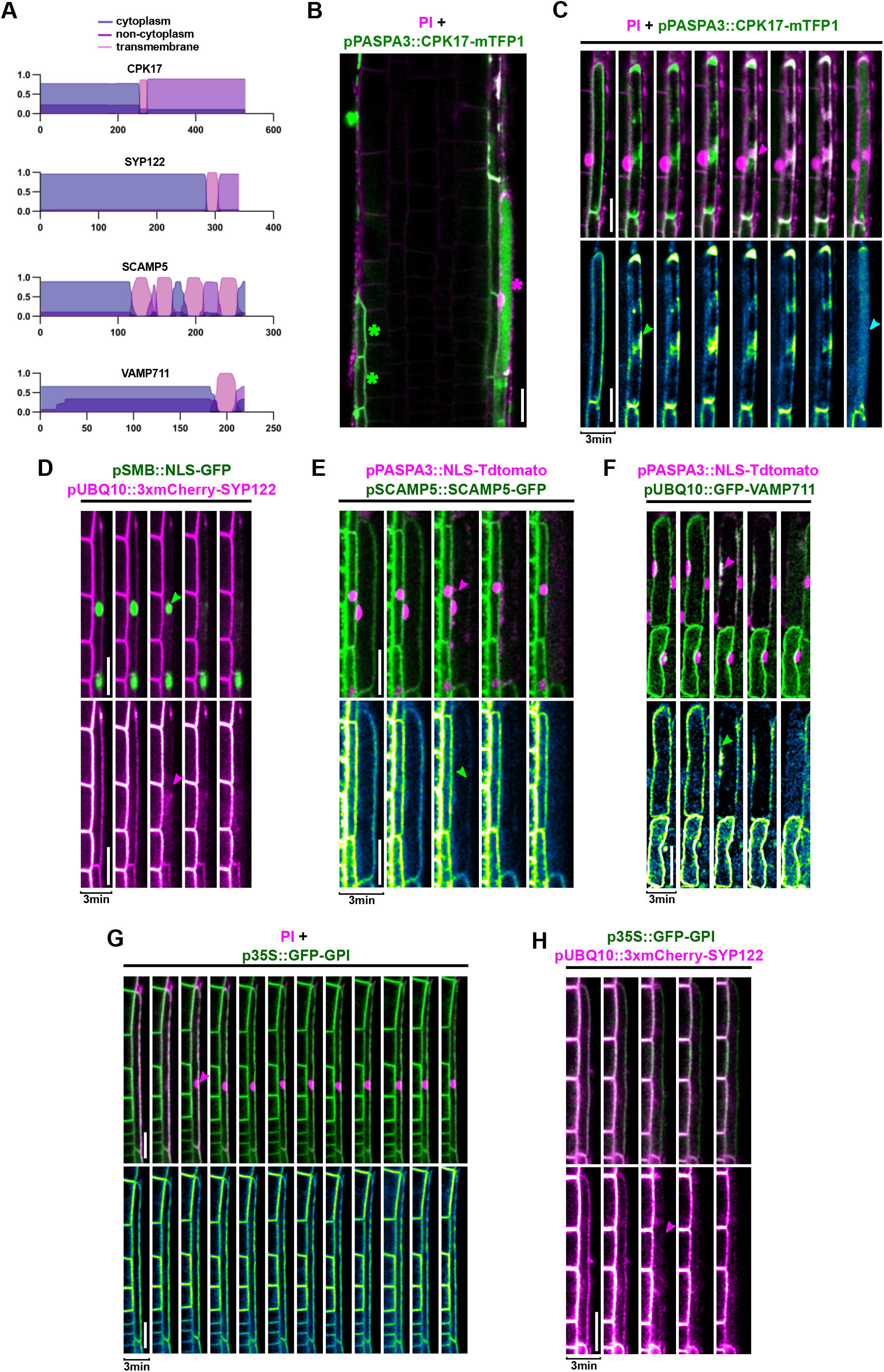
Dynamic behaviors of plasma membrane- and tonoplast-localized proteins during PCD execution. **(A)** Prediction of transmembrane domains in PM-localized (CPK17, SYP122 and SCAMP5) and tonoplast-localized (VAMP711) proteins. **(B)** Confocal images root cap cells expressing CPK17-mTFP1 in 4-day-old Arabidopsis seedlings. Green asterisks indicate living LRC cells with PM-localized CPK17-mTFP1, magenta asterisk indicate dying or dead cells. **(C)** Confocal time-lapse series of dying LRC cells. The upper panel shows PI in magenta and mTFP1 in green, the lower panel shows the mTFP1 channel separately. The green arrow indicates PM endodomain shedding, the magenta arrow indicates PI entry, and the cyan arrow indicates vacuolar collapse. **(D)** Confocal time-lapse series showing the simultaneous NE breakdown (green arrow) and PM shedding (magenta arrow) during PCD execution. The upper panel shows mCherry in magenta and NLS-GFP in green, the lower panel shows the mCherry channel separately. **(E)** Confocal time-lapse series showing the simultaneous NE breakdown (magenta arrow) and PM shedding (green arrow) during PCD execution. The upper panel shows NLS-TdTOMATO in magenta and SCAMP5-GFP in green, the lower panel shows the GFP channel separately. **(F)** Confocal time-lapse series showing the simultaneous NE breakdown (magenta arrow) and solubilization of the vacuole membrane marker GFP-VAMP711 (green arrow) during PCD execution. The upper panel shows NLS-TdTOMATO in magenta and GFP-VAMP711 in green, the lower panel shows the GFP channel separately. **(G)** Confocal time-lapse series showing PI entry (magenta arrow) and maintenance of apoplastic GFP-GPI after cell death execution. The upper panel shows PI in magenta and GFP-GPI in green, the lower panel shows the GFP channel separately. **(H)** Confocal time-lapse series of a dual marker line of mCherry-SYP122 (magenta, fluorescent tag on the cytoplasmic side) and GFP-GPI (green, fluorescent tag on the apoplastic side) showing the solubilization of mCherry in the cytoplasm (magenta arrow), while the extracellular GFP-GPI remains PM-bound. Scale bars are 20 μm.

**Supplemental Figure S2:**
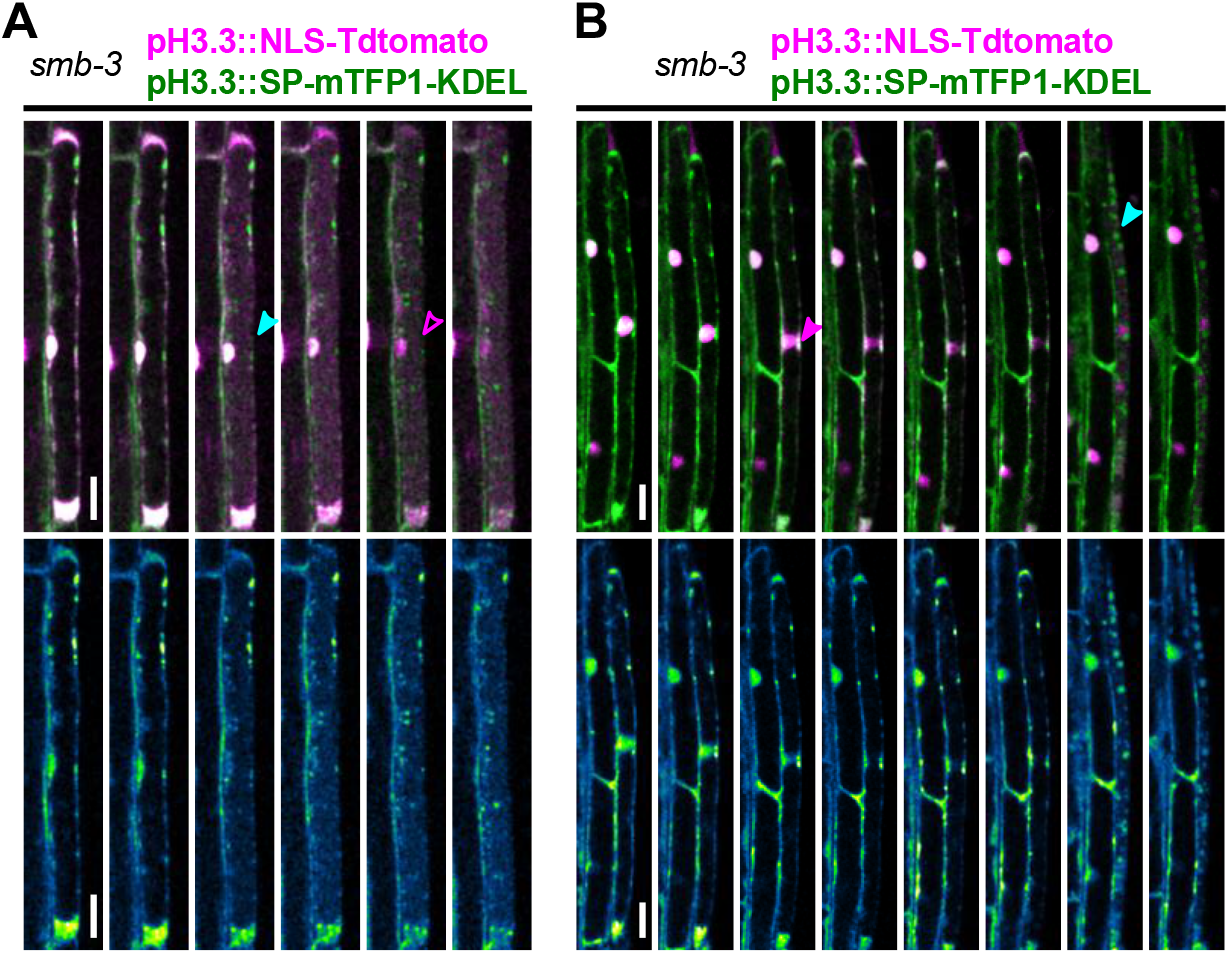
Aberrant dynamics of an ER lumen marker during cell death in the *smb-3* mutant. **(A-B)** Two representative confocal time-lapse series of dying LRC cells in 5-day old *smb-3* mutants. Upper panels show NLS-TdTOMATO in magenta and mTFP1 in green, the lower panels show the mTFP1 channel separately. There is no clearly defined release of the ER lumen marker *pH3.3::SP-mTFP1-KDEL* after vacuolar collapse (cyan arrow) and NE breakdown (magenta arrow). Scale bars are 20 μm.

**Supplemental Figure S3:**
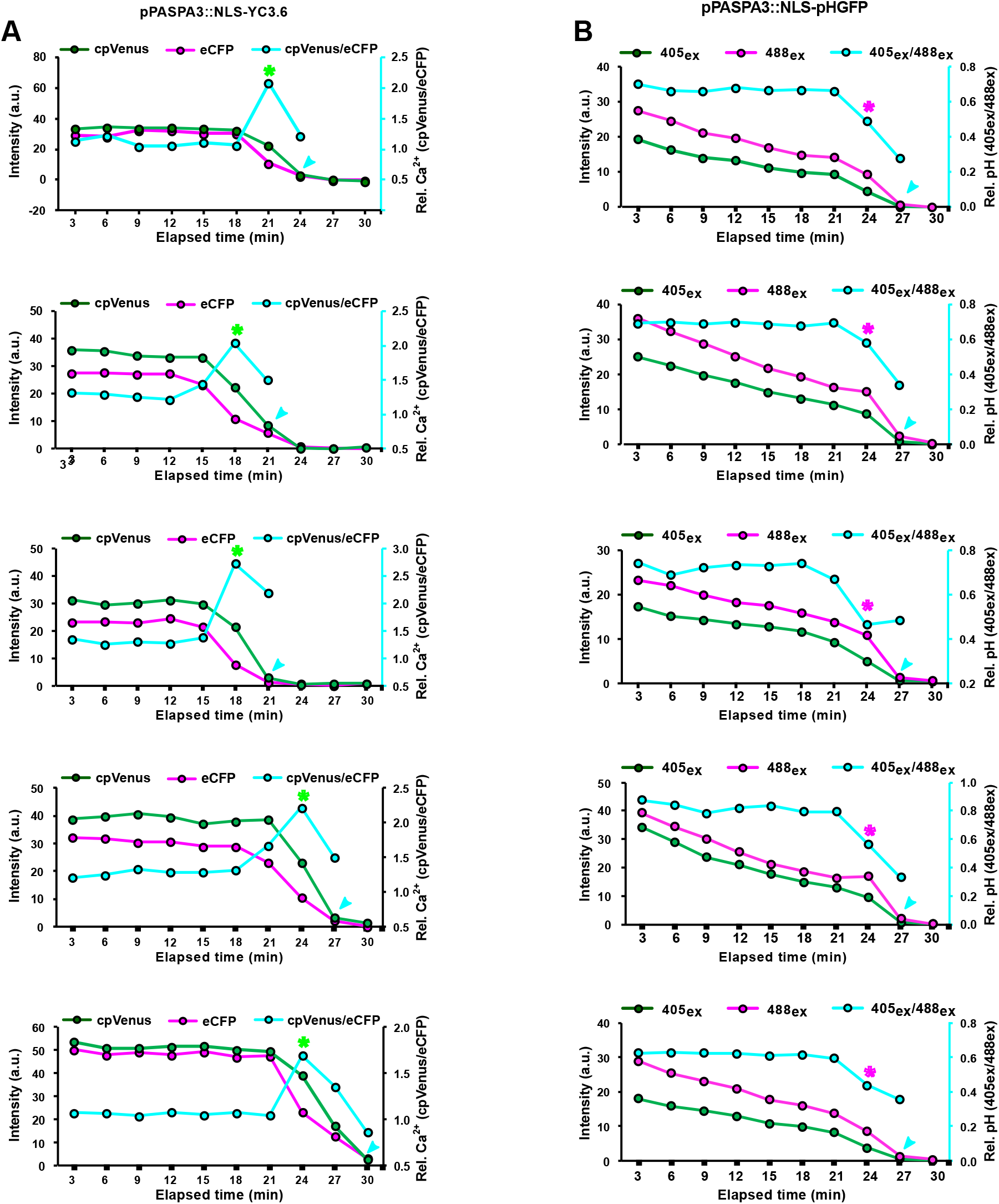
Intracellular calcium transient and acidification precede PCD execution. Five representative graphs related to figure 4C and D, showing the dynamics and ratios of fluorescent intensities correlating to intracellular [Ca^2+^] **(A)** and pH **(B)** sensors, respectively, expressing in wild-type LRC cells. The ratios of fluorescence intensity were obtained after background noise subtraction for each channel and corresponding to the relative [Ca^2+^]_nuc_ (cpVenus/eCFP) or pH_nuc_ (405ex/488ex). The cyan arrow indicates NE breakdown, the green asterisk in panel A marks the Ca^2+^ transient and the magenta asterisk in panel B marks acidification.

**Supplemental Figure S4:**
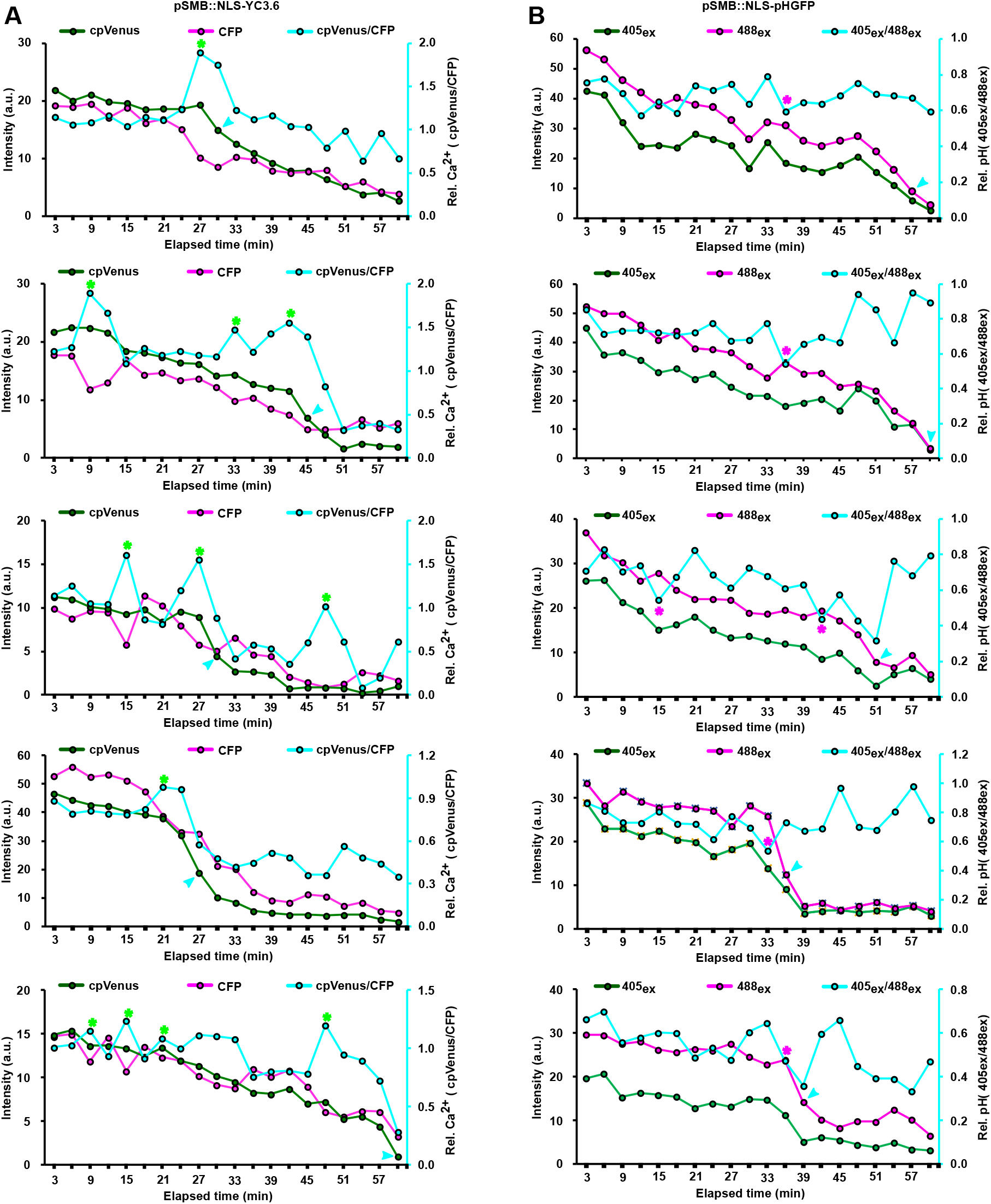
Aberrant pattern of intracellular calcium transient and acidification during *smb-3* cell death. Five representative graphs related to figure 5C and D, showing the dynamics and ratios of fluorescent intensities correlating to intracellular [Ca^2+^] **(A)** and pH **(B)** sensors, respectively, expressing in *smb-3* LRC cells. The ratios of fluorescence intensity were obtained after background noise subtraction for each channel and corresponding to the relative [Ca^2+^]_nuc_ (cpVenus/eCFP) or pH_nuc_ (405ex/488ex). The cyan arrow indicates NE breakdown, the green asterisk in panel A marks the Ca^2+^ transient and the magenta asterisk in panel B marks acidification.

**Supplemental Figure S5:**
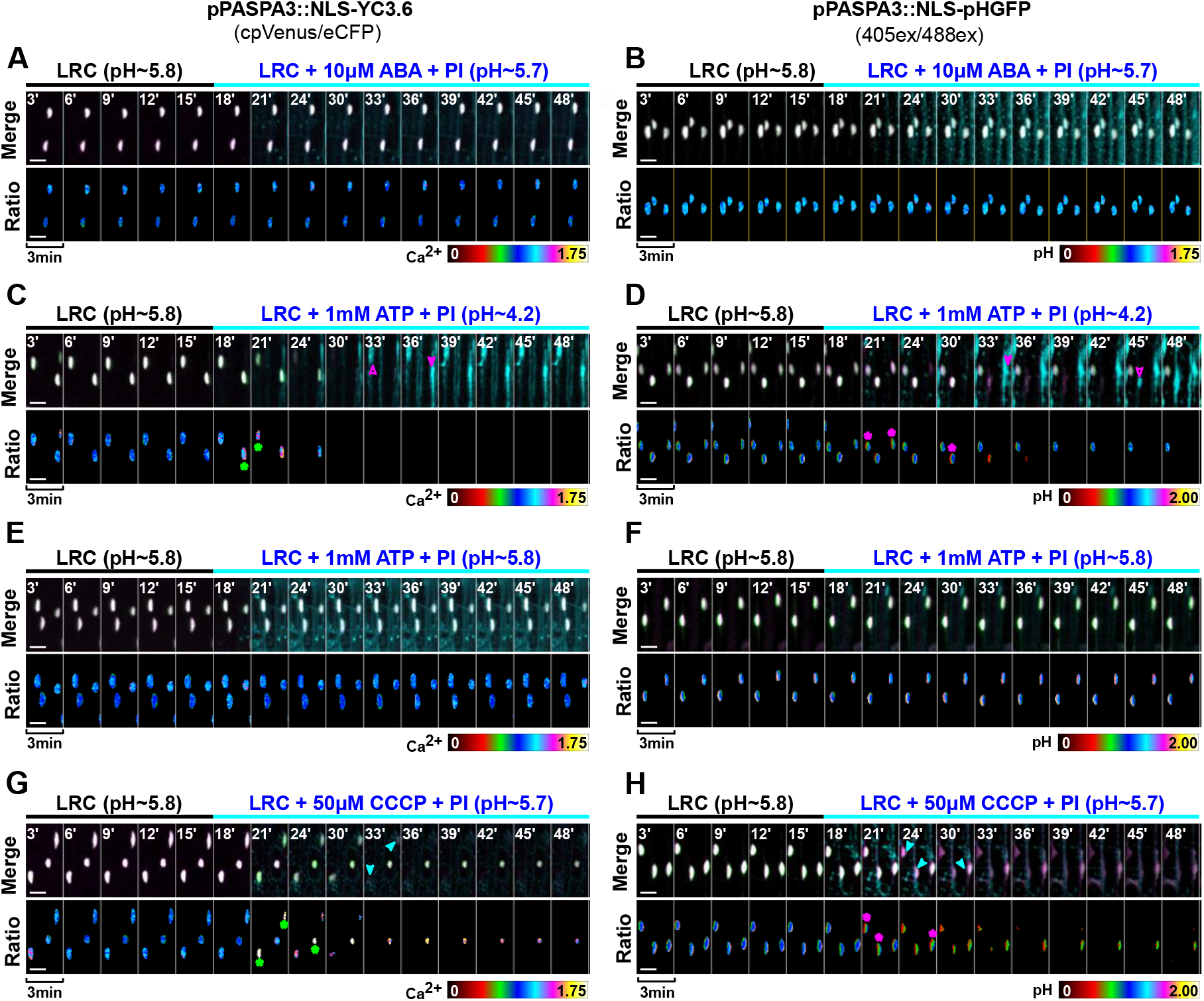
Pharmacological treatment causes distinct [Ca^2+^]_nuc_ and pH_nuc_ signatures, which are followed by cell death specifically in distal LRC cells. **(A,C,E,G)** Time-lapse confocal images of LRC cells from 4-day-old Col-0 seedlings expressing *pPASPA3::NLS-YC3.6* in response to ABA (n=4 roots), ATP (pH∼4.2, n=5 roots; pH∼5.8, n=4 roots), and CCCP (n=6 roots) application, respectively. Magenta arrows and cyan arrows indicate PI entry and NE breakdown, respectively, and green asterisks mark the [Ca^2+^]_nuc_ elevation. **(B,D,F,H)** Time-lapse confocal images of LRC cells from 4-day-old Col-0 seedlings expressing *pPASPA3::NLS-NLSpHGFP* in response to ABA (n=7 roots), ATP (pH∼4.2, n=4 roots; pH∼5.8, n=6 roots) and CCCP (n=6 roots) application, respectively. Magenta arrows and cyan arrows indicate PI entry and NE breakdown respectively, and magenta asterisks mark pH_nuc_ acidification. Upper panels show the merged signals, lower panels show ratiometric imaging. Scale bars are 10 μm.

**Supplemental Figure S6:**
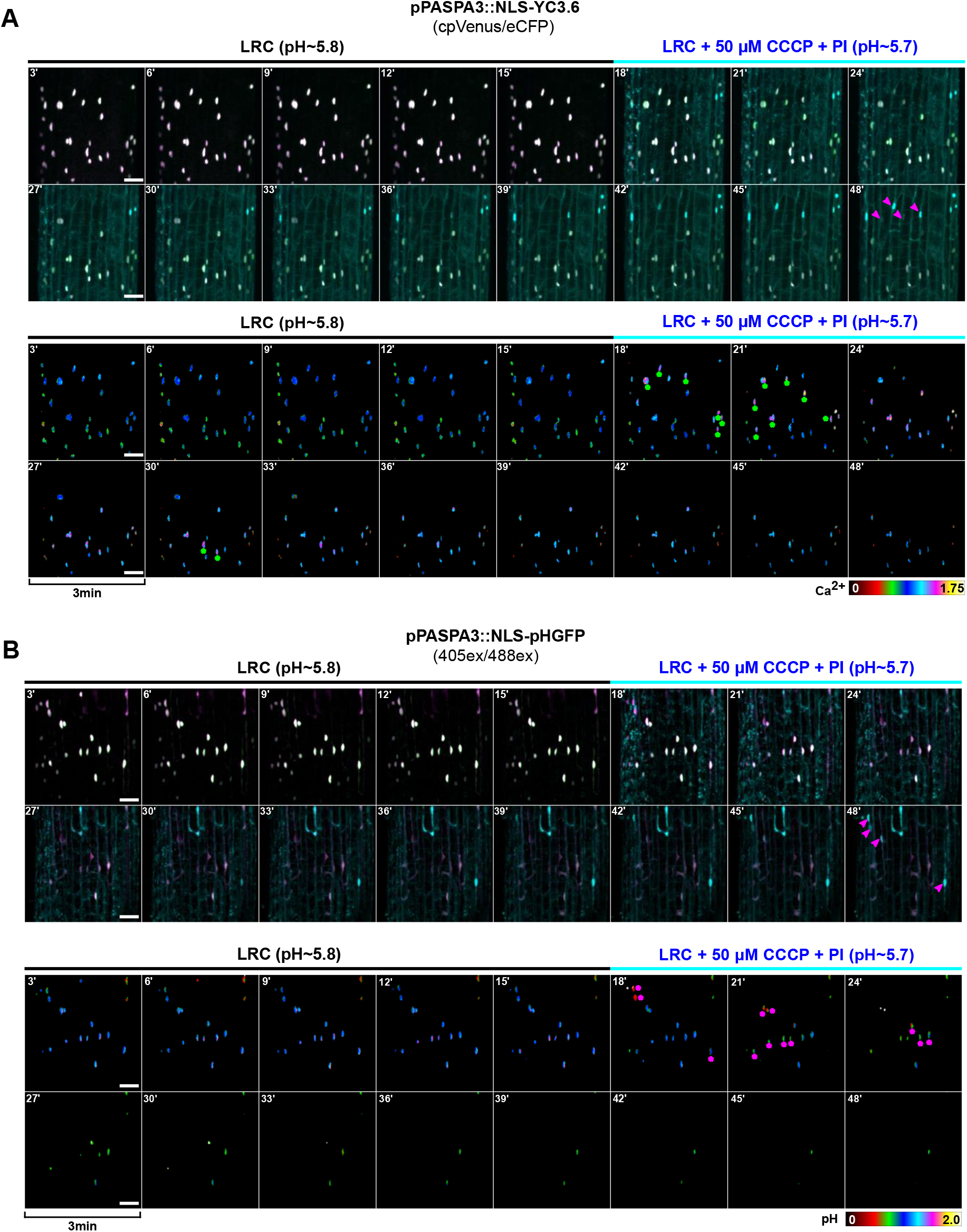
CCCP treatment causes [Ca^2+^]_nuc_ and pH_nuc_ signatures, which are followed by cell death specifically in distal LRC cells. **(A)** Time-lapse confocal images of LRC cells from 4-day-old Col-0 seedlings expressing *pPASPA3::NLS-YC3.6* in response to CCCP application. Magenta arrows indicate PI entry, and green asterisks mark the [Ca^2+^]_nuc_ elevation. **(B)** Time-lapse confocal images of LRC cells from 4-day-old Col-0 seedlings expressing *pPASPA3::NLS-NLSpHGFP* in response to CCCP application. Magenta arrows and cyan arrows indicate PI entry, and magenta asterisks mark pH_nuc_ acidification. Upper panels show the merged signals, lower panels show ratiometric imaging. Scale bars are 20 μm.

**Supplemental Figure S7:**
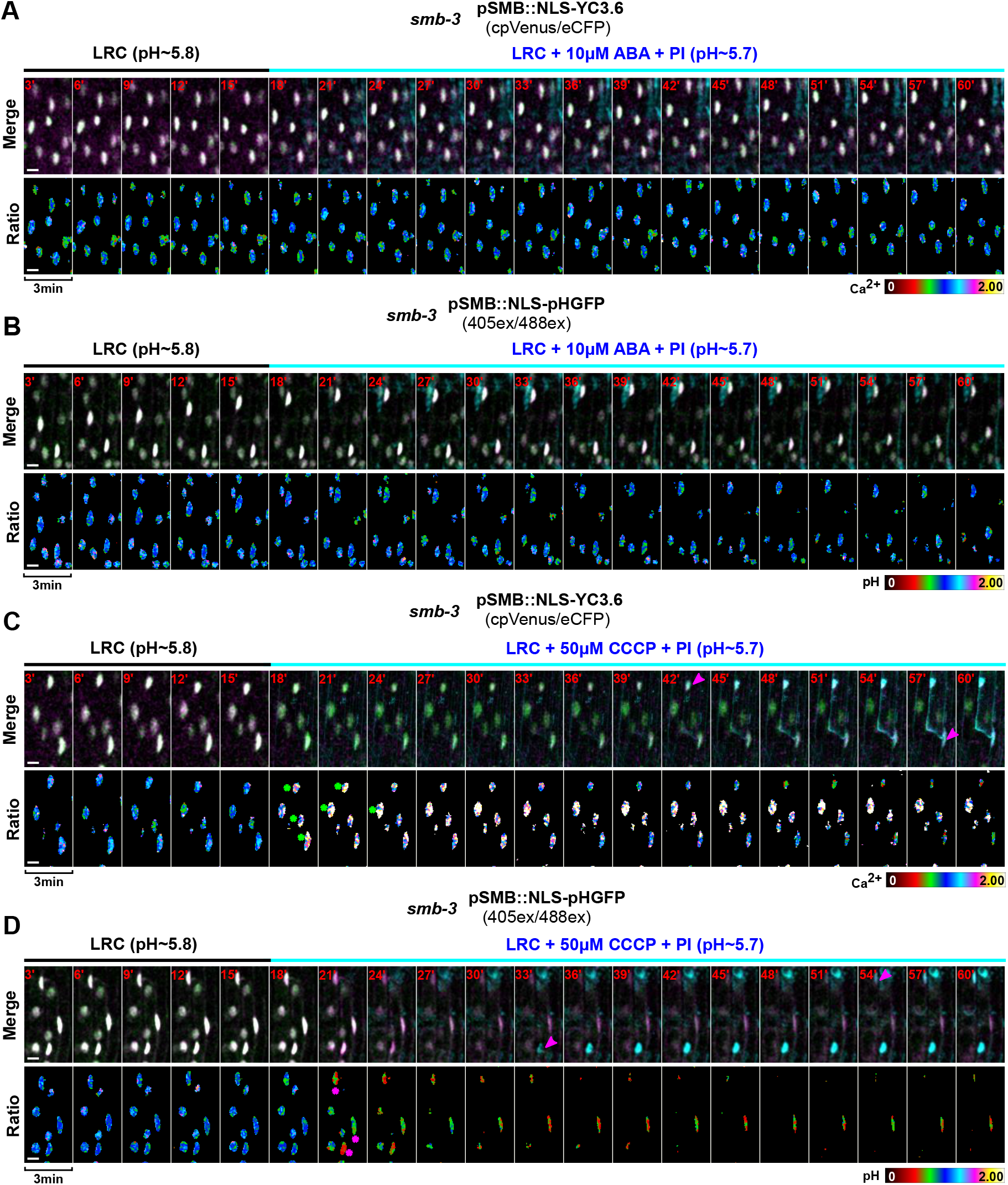
Short-term CCCP treatment induces intracellular calcium elevation and acidification in *smb-3* LRC cells. **(A,C)** Time-lapse confocal images of LRC cells from 5-day-old *smb-3* seedlings expressing *pSMB::NLS-YC3.6* showing the response to ABA (n=3 roots) and CCCP (n=7 roots) treatment, respectively. Magenta arrows indicate PI entry, and green asterisks mark the [Ca^2+^]_nuc_ elevation. **(B,D)** Time-lapse confocal images of LRC cells from 5-day-old *smb-3* seedlings expressing *pPASPA3::NLS-pHGFP* showing the response to ABA (n=4 roots) and CCCP (n=8 roots) treatment, respectively. Magenta arrows indicate PI entry, and magenta asterisks mark the pH_nuc_ acidification. Upper panels show the merged signals, lower panels show ratiometric imaging. Scale bars are 10 μm.

**Supplemental Table S1:**
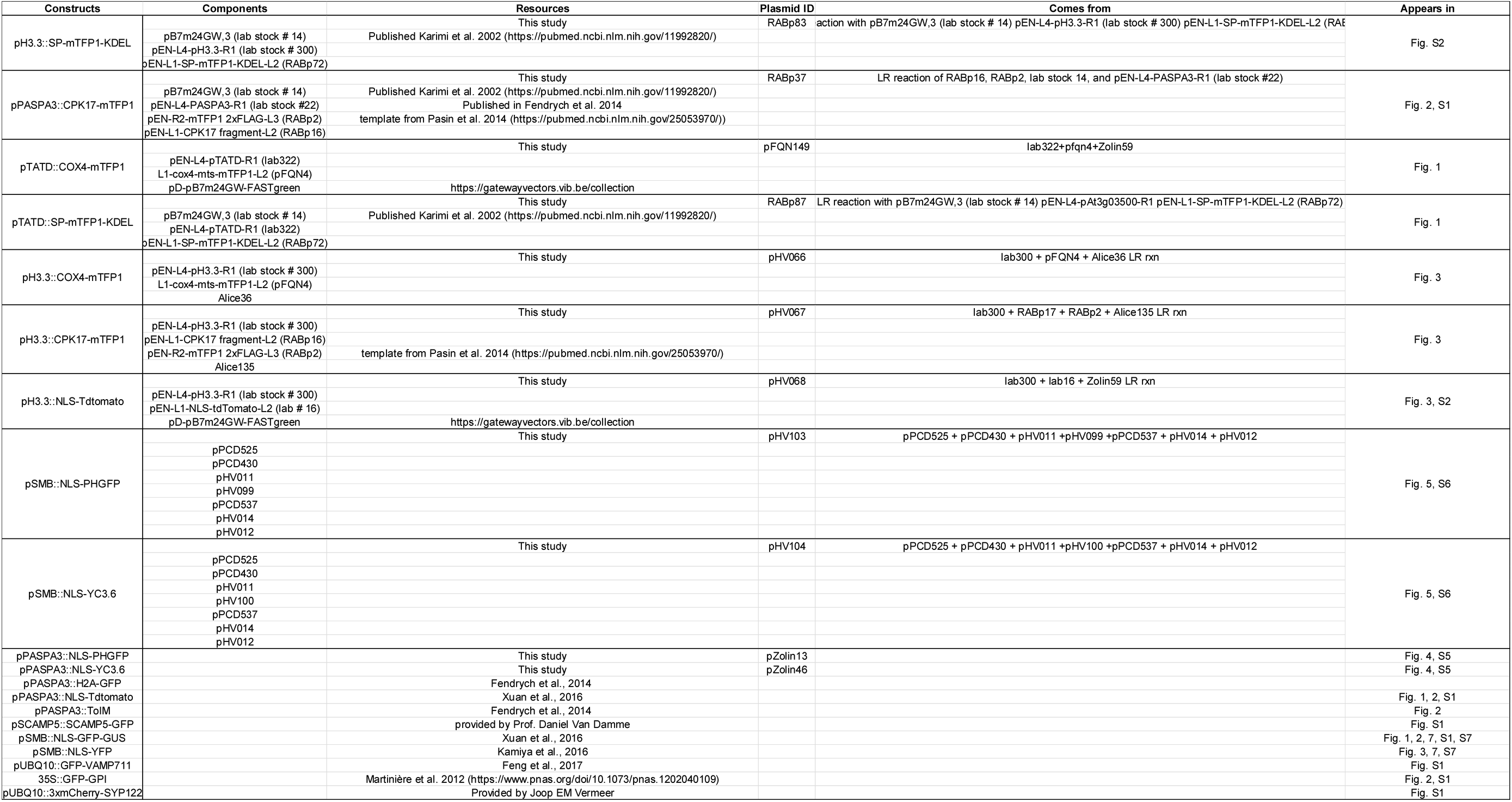
Plasmid information.

**Supplemental Table S2:**
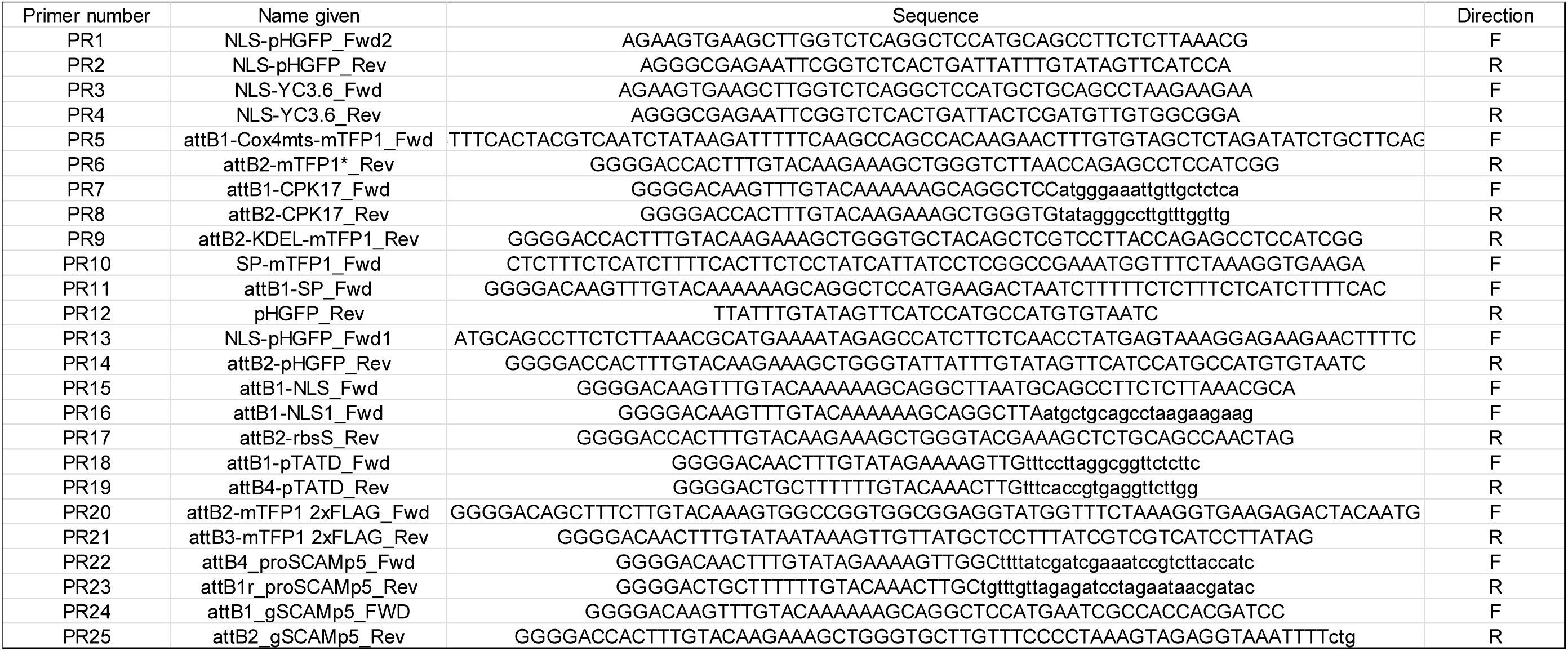
Primer information and sequences.

**Supplemental Table S3:**
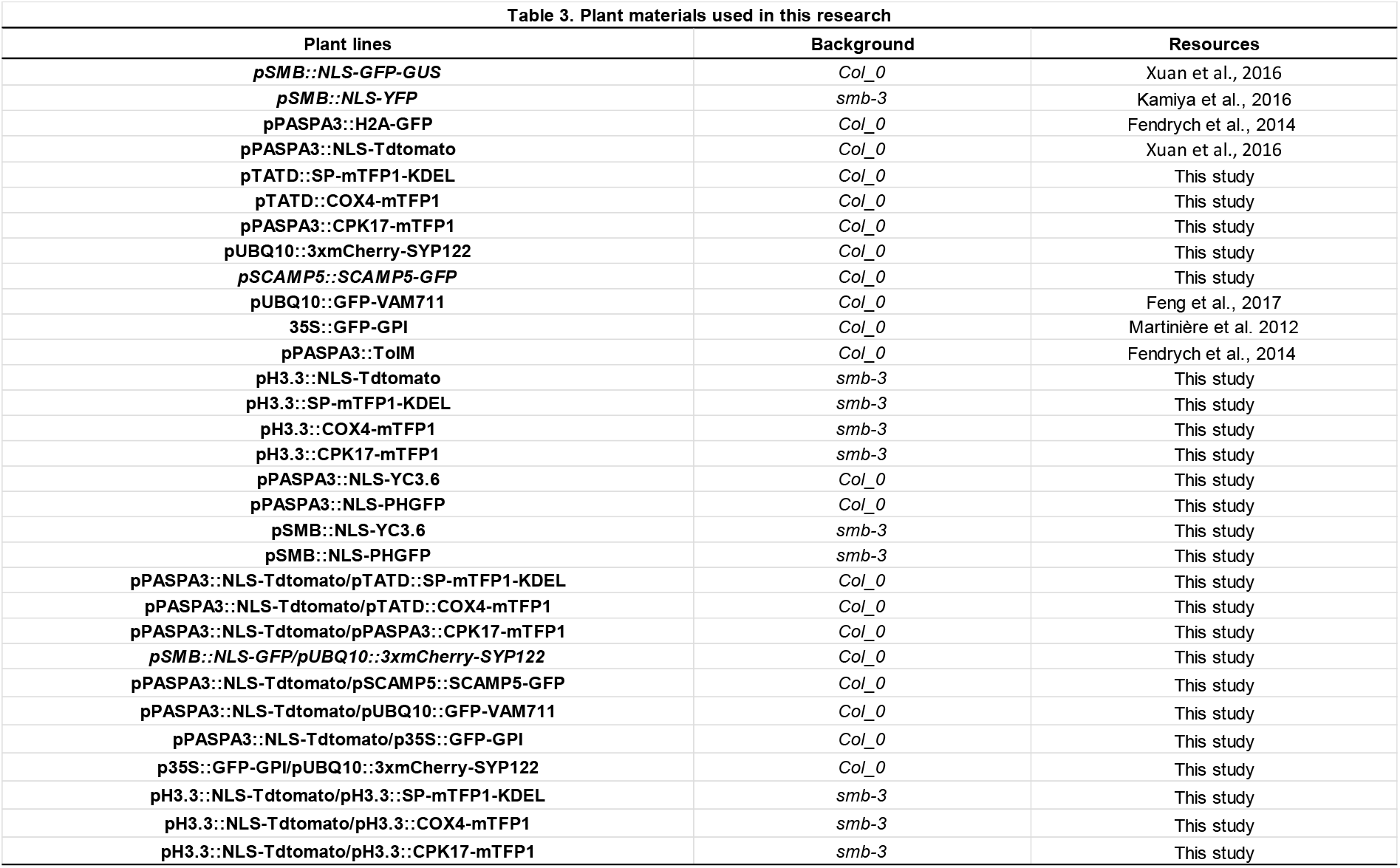
Information on transgenic plant lines used in this study.

| Table 3. Plant materials used in this research |  |  |
| --- | --- | --- |
| Plant lines | Background | Resources |
| *pSMB::NLS-GFP-GUS* | *Col_0* | Xuan et al., 2016 |
| *pSMB::NLS-YFP* | *smb-3* | Kamiya et al., 2016 |
| *pPASPA3::H2A-GFP* | *Col_0* | Fendrych et al., 2014 |
| *pPASPA3::NLS-Tdtomato* | *Col_0* | Xuan et al., 2016 |
| *pTATD::SP-mTFP1-KDEL* | *Col_0* | This study |
| *pTATD::COX4-mTFP1* | *Col_0* | This study |
| *pPASPA3::CPK17-mTFP1* | *Col_0* | This study |
| *pUBQ10::3xmCherry-SYP122* | *Col_0* | This study |
| *pSCAMP5::SCAMP5-GFP* | *Col_0* | This study |
| *pUBQ10::GFP-VAM711* | *Col_0* | Feng et al., 2017 |
| *35S::GFP-GPI* | *Col_0* | Martinière et al. 2012 |
| *pPASPA3::ToIM* | *Col_0* | Fendrych et al., 2014 |
| *pH3.3::NLS-Tdtomato* | *smb-3* | This study |
| *pH3.3::SP-mTFP1-KDEL* | *smb-3* | This study |
| *pH3.3::COX4-mTFP1* | *smb-3* | This study |
| *pH3.3::CPK17-mTFP1* | *smb-3* | This study |
| *pPASPA3::NLS-YC3.6* | *Col_0* | This study |
| *pPASPA3::NLS-PHGFP* | *Col_0* | This study |
| *pSMB::NLS-YC3.6* | *smb-3* | This study |
| *pSMB::NLS-PHGFP* | *smb-3* | This study |
| *pPASPA3::NLS-Tdtomato/pTATD::SP-mTFP1-KDEL* | *Col_0* | This study |
| *pPASPA3::NLS-Tdtomato/pTATD::COX4-mTFP1* | *Col_0* | This study |
| *pPASPA3::NLS-Tdtomato/pPASPA3::CPK17-mTFP1* | *Col_0* | This study |
| *pSMB::NLS-GFP/pUBQ10::3xmCherry-SYP122* | *Col_0* | This study |
| *pPASPA3::NLS-Tdtomato/pSCAMP5::SCAMP5-GFP* | *Col_0* | This study |
| *pPASPA3::NLS-Tdtomato/pUBQ10::GFP-VAM711* | *Col_0* | This study |
| *pPASPA3::NLS-Tdtomato/p35S::GFP-GPI* | *Col_0* | This study |
| *p35S::GFP-GPI/pUBQ10::3xmCherry-SYP122* | *Col_0* | This study |
| *pH3.3::NLS-Tdtomato/pH3.3::SP-mTFP1-KDEL* | *smb-3* | This study |
| *pH3.3::NLS-Tdtomato/pH3.3::COX4-mTFP1* | *smb-3* | This study |
| *pH3.3::NLS-Tdtomato/pH3.3::CPK17-mTFP1* | *smb-3* | This study |

## References

Andersen, T.G., Naseer, S., Ursache, R., Wybouw, B., Smet, W., De Rybel, B., Vermeer, J.E.M., and Geldner, N. (2018). Diffusible repression of cytokinin signalling produces endodermal symmetry and passage cells. Nature 555, 529–533.

Balk, J., and Leaver, C.J. (2001). The PET1-CMS mitochondrial mutation in sunflower is associated with premature programmed cell death and cytochrome c release. Plant Cell 13, 1803–1818.

Balk, J., Leaver, C.J., and McCabe, P.F. (1999). Translocation of cytochrome c from the mitochondria to the cytosol occurs during heat-induced programmed cell death in cucumber plants. FEBS Lett 463, 151–154.

Battistelli, M., and Falcieri, E. (2020). Apoptotic Bodies: Particular Extracellular Vesicles Involved in Intercellular Communication. Biology (Basel) 9.

Behera, S., Zhaolong, X., Luoni, L., Bonza, M.C., Doccula, F.G., De Michelis, M.I., Morris, R.J., Schwarzlander, M., and Costa, A. (2018). Cellular Ca (2+) Signals Generate Defined pH Signatures in Plants. Plant Cell 30, 2704–2719.

Benetka, W., Mehlmer, N., Maurer-Stroh, S., Sammer, M., Koranda, M., Neumuller, R., Betschinger, J., Knoblich, J.A., Teige, M., and Eisenhaber, F. (2008). Experimental testing of predicted myristoylation targets involved in asymmetric cell division and calcium-dependent signalling. Cell Cycle 7, 3709–3719.

Bennett, T., van den Toorn, A., Sanchez-Perez, G.F., Campilho, A., Willemsen, V., Snel, B., and Scheres, B. (2010). SOMBRERO, BEARSKIN1, and BEARSKIN2 regulate root cap maturation in Arabidopsis. Plant Cell 22, 640–654.

Choi, J., Tanaka, K., Cao, Y., Qi, Y., Qiu, J., Liang, Y., Lee, S.Y., and Stacey, G. (2014). Identification of a plant receptor for extracellular ATP. Science 343, 290–294.

Cubría-Radío, M., and Nowack, M.K. (2019). Chapter Seven - Transcriptional networks orchestrating programmed cell death during plant development. In Current Topics in Developmental Biology, U. Grossniklaus, ed (Academic Press), pp. 161–184.

Daneva, A., Gao, Z., Van Durme, M., and Nowack, M.K. (2016). Functions and Regulation of Programmed Cell Death in Plant Development. Annu Rev Cell Dev Biol 32, 441–468.

Decaestecker, W., Buono, R.A., Pfeiffer, M.L., Vangheluwe, N., Jourquin, J., Karimi, M., Van Isterdael, G., Beeckman, T., Nowack, M.K., and Jacobs, T.B. (2019). CRISPR-TSKO: A Technique for Efficient Mutagenesis in Specific Cell Types, Tissues, or Organs in Arabidopsis. Plant Cell 31, 2868–2887.

Doccula, F.G., Luoni, L., Behera, S., Bonza, M.C., and Costa, A. (2018). In Vivo Analysis of Calcium Levels and Glutathione Redox Status in Arabidopsis Epidermal Leaf Cells Infected with the Hypersensitive Response-Inducing Bacteria Pseudomonas syringae pv. tomato AvrB (PstAvrB). Methods Mol Biol 1743, 125–141.

Domínguez, F., and Cejudo, F.J. (2012). A comparison between nuclear dismantling during plant and animal programmed cell death. Plant Sci. 197, 114–121.

Escamez, S., and Tuominen, H. (2014). Programmes of cell death and autolysis in tracheary elements: when a suicidal cell arranges its own corpse removal. J Exp Bot 65, 1313–1321.

Farage-Barhom, S., Burd, S., Sonego, L., Mett, A., Belausov, E., Gidoni, D., and Lers, A. (2011). Localization of the Arabidopsis senescence- and cell death-associated BFN1 nuclease: from the ER to fragmented nuclei. Mol Plant 4, 1062–1073.

Fendrych, M., Van Hautegem, T., Van Durme, M., Olvera-Carrillo, Y., Huysmans, M., Karimi, M., Lippens, S., Guerin, C.J., Krebs, M., Schumacher, K., and Nowack, M.K. (2014). Programmed cell death controlled by ANAC033/SOMBRERO determines root cap organ size in Arabidopsis. Curr Biol 24, 931–940.

Feng, Q.N., Song, S.J., Yu, S.X., Wang, J.G., Li, S., and Zhang, Y. (2017). Adaptor Protein-3-Dependent Vacuolar Trafficking Involves a Subpopulation of COPII and HOPS Tethering Proteins. Plant Physiol 174, 1609–1620.

Fukuda, H. Programmed cell death of tracheary elements as a paradigm in plants.

Groover, A., and Jones, A.M. (1999). Tracheary element differentiation uses a novel mechanism coordinating programmed cell death and secondary cell wall synthesis. Plant Physiol 119, 375–384.

Hara-Nishimura, I., and Hatsugai, N. (2011). The role of vacuole in plant cell death. Cell Death Differ 18, 1298–1304.

Hierl, G., Howing, T., Isono, E., Lottspeich, F., and Gietl, C. (2014). Ex vivo processing for maturation of Arabidopsis KDEL-tailed cysteine endopeptidase 2 (AtCEP2) pro-enzyme and its storage in endoplasmic reticulum derived organelles. Plant Mol Biol 84, 605–620.

Huysmans, M., Lema, A.S., Coll, N.S., and Nowack, M.K. (2017). Dying two deaths - programmed cell death regulation in development and disease. Curr Opin Plant Biol 35, 37–44.

Huysmans, M., Buono, R.A., Skorzinski, N., Radio, M.C., De Winter, F., Parizot, B., Mertens, J., Karimi, M., Fendrych, M., and Nowack, M.K. (2018). NAC Transcription Factors ANAC087 and ANAC046 Control Distinct Aspects of Programmed Cell Death in the Arabidopsis Columella and Lateral Root Cap. Plant Cell 30, 2197–2213.

Ingouff, M., Selles, B., Michaud, C., Vu, T.M., Berger, F., Schorn, A.J., Autran, D., Van Durme, M., Nowack, M.K., Martienssen, R.A., and Grimanelli, D. (2017). Live-cell analysis of DNA methylation during sexual reproduction in Arabidopsis reveals context and sex-specific dynamics controlled by noncanonical RdDM. Genes Dev 31, 72–83.

Jiang, C., Wang, J., Leng, H.N., Wang, X., Liu, Y., Lu, H., Lu, M.Z., and Zhang, J. (2021). Transcriptional Regulation and Signaling of Developmental Programmed Cell Death in Plants. Front Plant Sci 12, 702928.

Jones, A.M. (2001). Programmed cell death in development and defense. Plant Physiol 125, 94–97.

Kabbage, M., Kessens, R., Bartholomay, L.C., and Williams, B. (2017). The Life and Death of a Plant Cell. Annu Rev Plant Biol 68, 375–404.

Kamiya, M., Higashio, S.Y., Isomoto, A., Kim, J.M., Seki, M., Miyashima, S., and Nakajima, K. (2016). Control of root cap maturation and cell detachment by BEARSKIN transcription factors in Arabidopsis. Development 143, 4063–4072.

Karimi, M., Inze, D., and Depicker, A. (2002). GATEWAY vectors for Agrobacterium-mediated plant transformation. Trends Plant Sci 7, 193–195.

Karimi, M., Depicker, A., and Hilson, P. (2007). Recombinational cloning with plant gateway vectors. Plant Physiol 145, 1144–1154.

Krebs, M., and Schumacher, K. (2013). Live cell imaging of cytoplasmic and nuclear Ca2+ dynamics in Arabidopsis roots. Cold Spring Harb Protoc 2013, 776–780.

Krebs, M., Held, K., Binder, A., Hashimoto, K., Den Herder, G., Parniske, M., Kudla, J., and Schumacher, K. (2012). FRET-based genetically encoded sensors allow high-resolution live cell imaging of Ca (2) (+) dynamics. Plant J 69, 181–192.

Kumpf, R.P., and Nowack, M.K. (2015). The root cap: a short story of life and death. J Exp Bot 66, 5651–5662.

Lampropoulos, A., Sutikovic, Z., Wenzl, C., Maegele, I., Lohmann, J.U., and Forner, J. (2013). GreenGate---a novel, versatile, and efficient cloning system for plant transgenesis. PLoS ONE 8, e83043.

Lin, Z., Xie, F., Trivino, M., Karimi, M., Bosch, M., Franklin-Tong, V.E., and Nowack, M.K. (2020). Ectopic Expression of a Self-Incompatibility Module Triggers Growth Arrest and Cell Death in Vegetative Cells. Plant Physiol 183, 1765–1779.

Logan, D.C. (2006). The mitochondrial compartment. J Exp Bot 57, 1225–1243.

Martiniere, A., Lavagi, I., Nageswaran, G., Rolfe, D.J., Maneta-Peyret, L., Luu, D.T., Botchway, S.W., Webb, S.E., Mongrand, S., Maurel, C., Martin-Fernandez, M.L., Kleine-Vehn, J., Friml, J., Moreau, P., and Runions, J. (2012). Cell wall constrains lateral diffusion of plant plasma-membrane proteins. Proc Natl Acad Sci U S A 109, 12805–12810.

McArthur, K., Whitehead, L.W., Heddleston, J.M., Li, L., Padman, B.S., Oorschot, V., Geoghegan, N.D., Chappaz, S., Davidson, S., San Chin, H., Lane, R.M., Dramicanin, M., Saunders, T.L., Sugiana, C., Lessene, R., Osellame, L.D., Chew, T.-L., Dewson, G., Lazarou, M., Ramm, G., Lessene, G., Ryan, M.T., Rogers, K.L., van Delft, M.F., and Kile, B.T. (2018). BAK/BAX macropores facilitate mitochondrial herniation and mtDNA efflux during apoptosis. Science 359, eaao6047.

Mehlmer, N., Parvin, N., Hurst, C.H., Knight, M.R., Teige, M., and Vothknecht, U.C. (2012). A toolset of aequorin expression vectors for in planta studies of subcellular calcium concentrations in Arabidopsis thaliana. J Exp Bot 63, 1751–1761.

Minina, E.A., Dauphinee, A.N., Ballhaus, F., Gogvadze, V., Smertenko, A.P., and Bozhkov, P.V. (2021). Apoptosis is not conserved in plants as revealed by critical examination of a model for plant apoptosis-like cell death. BMC Biol 19, 100.

Moseyko, N., and Feldman, L.J. (2001). Expression of pH-sensitive green fluorescent protein in Arabidopsis thaliana. Plant Cell Environ 24, 557–563.

Nagai, T., Yamada, S., Tominaga, T., Ichikawa, M., and Miyawaki, A. (2004). Expanded dynamic range of fluorescent indicators for Ca (2+) by circularly permuted yellow fluorescent proteins. Proc Natl Acad Sci U S A 101, 10554–10559.

Nelson, B.K., Cai, X., and Nebenfuhr, A. (2007). A multicolored set of in vivo organelle markers for co-localization studies in Arabidopsis and other plants. Plant J 51, 1126–1136.

Obara, K., Kuriyama, H., and Fukuda, H. (2001). Direct evidence of active and rapid nuclear degradation triggered by vacuole rupture during programmed cell death in Zinnia. Plant Physiol 125, 615–626.

Olvera-Carrillo, Y., Van Bel, M., Van Hautegem, T., Fendrych, M., Huysmans, M., Simaskova, M., van Durme, M., Buscaill, P., Rivas, S., Coll, N.S., Coppens, F., Maere, S., and Nowack, M.K. (2015). A Conserved Core of Programmed Cell Death Indicator Genes Discriminates Developmentally and Environmentally Induced Programmed Cell Death in Plants. Plant Physiol 169, 2684–2699.

Pasin, F., Kulasekaran, S., Natale, P., Simon-Mateo, C., and Garcia, J.A. (2014). Rapid fluorescent reporter quantification by leaf disc analysis and its application in plant-virus studies. Plant Methods 10, 22.

Ren, H., Zhao, X., Li, W., Hussain, J., Qi, G., and Liu, S. (2021). Calcium Signaling in Plant Programmed Cell Death. Cells 10.

Rieger, A.M., Hall, B.E., Luong, L.T., Schang, L.M., and Barreda, D.R. (2010). Conventional apoptosis assays using propidium iodide generate a significant number of false positives that prevent accurate assessment of cell death. J. Immunol. Methods 358, 81–92.

Riley, J.S., Quarato, G., Cloix, C., Lopez, J., O’Prey, J., Pearson, M., Chapman, J., Sesaki, H., Carlin, L.M., Passos, J.F., Wheeler, A.P., Oberst, A., Ryan, K.M., and Tait, S.W. (2018). Mitochondrial inner membrane permeabilisation enables mtDNA release during apoptosis. EMBO J 37.

Santavanond, J.P., Rutter, S.F., Atkin-Smith, G.K., and Poon, I.K.H. (2021). Apoptotic Bodies: Mechanism of Formation, Isolation and Functional Relevance. Subcell Biochem 97, 61–88.

Schindelin, J., Arganda-Carreras, I., Frise, E., Kaynig, V., Longair, M., Pietzsch, T., Preibisch, S., Rueden, C., Saalfeld, S., Schmid, B., Tinevez, J.Y., White, D.J., Hartenstein, V., Eliceiri, K., Tomancak, P., and Cardona, A. (2012). Fiji: an open-source platform for biological-image analysis. Nat Methods 9, 676–682.

Tait, S.W., and Green, D.R. (2013). Mitochondrial regulation of cell death. Cold Spring Harb Perspect Biol 5.

Thomas, S.G., and Franklin-Tong, V.E. (2004). Self-incompatibility triggers programmed cell death in Papaver pollen. Nature 429, 305–309.

Toyooka, K., Goto, Y., Hashimoto, K., Wakazaki, M., Sato, M., and Hirai, M.Y. (2023). Endoplasmic Reticulum Bodies in the Lateral Root Cap are Involved in the Direct Transport of Beta-Glucosidase to Vacuoles. Plant and Cell Physiology, pcac177.

Truernit, E., and Haseloff, J. (2008). A simple way to identify non-viable cells within living plant tissue using confocal microscopy. Plant Methods 4, 15.

van Doorn, W.G., Beers, E.P., Dangl, J.L., Franklin-Tong, V.E., Gallois, P., Hara-Nishimura, I., Jones, A.M., Kawai-Yamada, M., Lam, E., Mundy, J., Mur, L.A., Petersen, M., Smertenko, A., Taliansky, M., Van Breusegem, F., Wolpert, T., Woltering, E., Zhivotovsky, B., and Bozhkov, P.V. (2011). Morphological classification of plant cell deaths. Cell Death Differ 18, 1241–1246.

Van Durme, M., and Nowack, M.K. (2016). Mechanisms of developmentally controlled cell death in plants. Curr Opin Plant Biol 29, 29–37.

Waadt, R., Koster, P., Andres, Z., Waadt, C., Bradamante, G., Lampou, K., Kudla, J., and Schumacher, K. (2020). Dual-Reporting Transcriptionally Linked Genetically Encoded Fluorescent Indicators Resolve the Spatiotemporal Coordination of Cytosolic Abscisic Acid and Second Messenger Dynamics in Arabidopsis. Plant Cell 32, 2582–2601.

Wang, L., Trivino, M., Lin, Z., Carli, J., Eaves, D.J., Van Damme, D., Nowack, M.K., Franklin-Tong, V.E., and Bosch, M. (2020). New opportunities and insights into Papaver self-incompatibility by imaging engineered Arabidopsis pollen. J Exp Bot 71, 2451–2463.

Wilkins, K.A., Bosch, M., Haque, T., Teng, N., Poulter, N.S., and Franklin-Tong, V.E. (2015). Self-incompatibility-induced programmed cell death in field poppy pollen involves dramatic acidification of the incompatible pollen tube cytosol. Plant Physiol 167, 766–779.

Willemsen, V., Bauch, M., Bennett, T., Campilho, A., Wolkenfelt, H., Xu, J., Haseloff, J., and Scheres, B. (2008). The NAC domain transcription factors FEZ and SOMBRERO control the orientation of cell division plane in Arabidopsis root stem cells. Dev Cell 15, 913–922.

Wu, J., Wang, S., Gu, Y., Zhang, S., Publicover, S.J., and Franklin-Tong, V.E. (2011). Self-incompatibility in Papaver rhoeas activates nonspecific cation conductance permeable to Ca2+ and K+. Plant Physiol 155, 963–973.

Xuan, W., Band, L.R., Kumpf, R.P., Van Damme, D., Parizot, B., De Rop, G., Opdenacker, D., Moller, B.K., Skorzinski, N., Njo, M.F., De Rybel, B., Audenaert, D., Nowack, M.K., Vanneste, S., and Beeckman, T. (2016). Cyclic programmed cell death stimulates hormone signaling and root development in Arabidopsis. Science 351, 384–387.

Yamaguchi, H., Maruyama, T., Urade, Y., and Nagata, S. (2014). Immunosuppression via adenosine receptor activation by adenosine monophosphate released from apoptotic cells. eLife 3, e02172.

Young, B., Wightman, R., Blanvillain, R., Purcel, S.B., and Gallois, P. (2010). pH-sensitivity of YFP provides an intracellular indicator of programmed cell death. Plant Methods 6, 27.

Yperman, K., Papageorgiou, A.C., Merceron, R., De Munck, S., Bloch, Y., Eeckhout, D., Jiang, Q., Tack, P., Grigoryan, R., Evangelidis, T., Van Leene, J., Vincze, L., Vandenabeele, P., Vanhaecke, F., Potocky, M., De Jaeger, G., Savvides, S.N., Tripsianes, K., Pleskot, R., and Van Damme, D. (2021). Distinct EH domains of the endocytic TPLATE complex confer lipid and protein binding. Nat Commun 12, 3050.

Yu, X.H., Perdue, T.D., Heimer, Y.M., and Jones, A.M. (2002). Mitochondrial involvement in tracheary element programmed cell death. Cell Death Differ 9, 189–198.

Zhang, D., Liu, D., Lv, X., Wang, Y., Xun, Z., Liu, Z., Li, F., and Lu, H. (2014). The cysteine protease CEP1, a key executor involved in tapetal programmed cell death, regulates pollen development in Arabidopsis. Plant Cell 26, 2939–2961.

Zhou, L.Z., Howing, T., Muller, B., Hammes, U.Z., Gietl, C., and Dresselhaus, T. (2016). Expression analysis of KDEL-CysEPs programmed cell death markers during reproduction in Arabidopsis. Plant Reprod 29, 265–272.

